# An intrinsic neuronal manifold underlies brain-wide hierarchical organization of behavior in *C. elegans*

**DOI:** 10.1101/2025.03.09.642241

**Authors:** Charles Fieseler, Itamar Lev, Ulises Rey, Lukas Hille, Hannah Brenner, Manuel Zimmer

## Abstract

Large-scale neuronal recordings in various species have revealed behavior-related activity patterns. Surprisingly, these activities are distributed across brain regions, including sensory areas. However, their origins and functions are incompletely understood. Using whole-brain imaging in freely behaving *C. elegans*, we discover that such distributed brain activity is dominated by a low dimensional manifold, corresponding to the major action sequence of *C. elegans*. The manifold originates largely from intrinsic dynamics, with minor contributions from movement-induced sensations. We discover a new function of the manifold: to gate the activity of a large number of neuron classes. This gating constrains them to selectively encode faster time-scale motor patterns during specific brain states. We therefore propose that one principle function of brain-wide dynamics is to enable hierarchical organization of behavior: information about major actions is broadcast via a distributed manifold, to enable nesting of granular motor patterns within those major actions.

## INTRODUCTION

Recent studies have discovered neural activity correlates of instantaneous behavior distributed throughout the brain, surprisingly, including sensory areas. These observations were made across animal phyla from invertebrates to mammals^1–6^, including primate cortical areas outside motor cortex, although to a significantly lesser extent^7,8^. The origins and functions of these activity patterns are not known^9^. Neuronal population activity patterns within specific brain regions, however, are well described. Such activity patterns are often modeled as smooth trajectories in a low dimensional neuronal space, termed a manifold^10–21^. For example, in primate motor cortical areas, both the planning and execution of arm reaches have been functionally linked to neuronal trajectories on low dimensional manifolds, demonstrating that manifolds can encode relevant aspects of behavior^10^. However, it is not known whether any manifold structure underlies the assembly of individual movements into long multi-action sequences. Moreover, it is unclear how neuronal activity underlying granular behaviors, such as grasping^22^, is nested within activity related to coarse behaviors such as arm reach.

We therefore hypothesized that one function of distributed brain-wide activity is to organize patterns of behavior across these different scales. In natural contexts, animals must simultaneously perform many tasks, from processing sensory information to coordinating actions and motor patterns, while maintaining longer-term behavioral goals. Ethological studies have proposed that this is achieved through a hierarchical organization of behavior, in which slower time-scale behaviors hierarchically orchestrate faster time-scale behaviors^23–27^. Some studies have identified neural activity correlates of hierarchical behavioral organization^28–31^ in specific neurons or brain regions. However, how such hierarchical organization is achieved across the brain and the role of neuronal manifolds in such organization, if any, is unknown.

Behavior-related brain activity may be generated solely internally or alternatively may originate from the sensory consequences of that behavior. For example, movement changes the visual landscape in the opposite direction of the movement, termed optical flow^32–34^. In many instances, such self-generation sensory information is suppressed in the nervous system^35^. Interestingly, multiple species also use it to adjust their movements, e.g. landing in flying insects^36^ and navigation in mammals including humans^37–39^. Additional sensory modalities may also exhibit self-generated signals. For example, when nematodes crawl backwards they sense traces of O_2_ and CO_2_ gases caused by their own metabolism^40^. Overall, such sensory-driven activity is highly correlated to the animals behavior and may be prevalent in the nervous system. Thus, disentangling the origins of brain-wide behavior-related activity is required for a comprehensive understanding of its functions.

The cross species findings of distributed behavior-related activity patterns and low dimensional manifolds suggest some universal principles^9^. In order to reveal these principles a set of open questions should be addressed: What are the origins and functions of brain-wide behavioral representations? Does any low dimensional structure both extend across the brain and serve behavioral functions, such as the establishment of behavioral hierarchies?

Despite considerable behavioral diversity, conserved principles can be well studied in tractable invertebrate models. The nematode *C. elegans* has a compact nervous system amenable to single cell resolution whole brain Ca^2+^ imaging of neuronal activity^41–43^. Recent work extended this imaging modality to freely moving animals^44–47^. Its locomotion behavior is characterized by a major action sequence of motor programs, each consisting of dorsal-ventral undulatory body waves. The motor programs are forward crawling (forward), backward crawling (reversal), and two types of turns in either direction of the dorsal-ventral body axis. After turning animals resume forward crawling, completing an action cycle^1,48–50^. Importantly, the timing of switching between actions appears spontaneously i.e., in the absence of acute sensory triggers^50^. Neuronal Ca^2+^ imaging in immobilized animals revealed population dynamics modeled by manifold trajectories that reflected spontaneous brain state progression through this major action sequence, in the absence of its motor execution^1^. Importantly, *C. elegans* has a stereotypic nervous system with identifiable cell classes across individuals, allowing direct comparison between immobilized and freely moving animals, as well as across mutant lines. Therefore this system allows the disambiguation of the intrinsic or extrinsic origins of the manifold. Furthermore, previous work described subtle head motor patterns as hierarchically nested within this major action sequence, and identified specific examples of hierarchically organized neuronal activity^28,31^. However, it remained unclear if brain-wide activity of freely moving animals (1) can be modeled as a manifold, (2) is dominated by intrinsic or extrinsic activity, and (3) displays hierarchical organization as a general principle.

In the present study we aimed to investigate the existence, origins, and functions of a neuronal manifold in freely moving animals. We developed a new pipeline for recording and analyzing brain-wide activity of freely moving animals. First, we reveal dominant and widely distributed brain-wide neuronal population activity that can be modeled as a low-dimension manifold and encodes the worm’s action sequence. We discovered that several oxygen and carbon dioxide sensory neuron classes contribute to the manifold by responding to re-afferent, i.e. movement induced, sensory stimuli. Crucially, the vast majority of shared population activity that forms the manifold can be interpreted as intrinsic, as it is present regardless of motor execution or structured sensory stimulation. Moreover, we find that many neuron classes that participate in the manifold exhibit additional higher-dimensional activity patterns in freely moving animals. In many neuron classes, this additional activity corresponds to granular faster time scale motor patterns. Using a Bayesian approach, we provide evidence supporting a model that the manifold, representing the upper-level action sequence, hierarchically gates residual activity corresponding to lower-level motor patterns. We propose that top-down hierarchical gating via a low-dimensional manifold is an organizational principle of the brain, used to establish behavioral motor hierarchies.

## RESULTS

### A pipeline to record and quantify whole brain activity and behavior in freely crawling *C. elegans*

In order to study the real-time relationship between neuronal activity and behavior, we devised a pipeline to record whole brain neuronal Ca^2+^ fluorescence imaging in freely crawling *C. elegans*. We therefore designed an assay arena where worms freely explore bare agar pads under a cover-glass. Brain activity was imaged with high magnification immersion optics, and behavior with infrared illumination and low magnification to capture the entire body of single worms. For this, we optimized a spinning disk confocal microscope and worm tracking stage^1,51^ (Fig. S1A). We used transgenic worms with GCaMP7b as a fluorescent Ca^2+^-indicator^52^ and mScarlet as a reference signal, both expressed in a pan-neuronal and nuclear-localized fashion^41^. We further generated a control strain with GFP instead of GCaMP, in order to calculate a baseline of motion-related artifacts.

To overcome the challenge of tracking the small and densely packed neurons in the deforming tissue of crawling worms, we took a segment-then-track strategy^53,54^. We first segmented the cellular nuclei using a custom trained 3D Stardist network^55,56^ and then tracked segmented objects across time^57^, using two trained SuperGlue neural networks^58^ (See Methods; Fig. S1B). We built a graphical user interface based on the Napari^59^ python package for manual correction of tracks, visualization of traces, and tools to facilitate neuronal identification. Using this graphical interface we produced six proof-read ground-truth data sets (four were used for training the tracking neural network, and two to quantify accuracy). Overall we estimate that median accuracy of tracked time points across neurons to be over 95% (Fig. S1C).

We identified a subset of neuronal identities based on their anatomy, position, and activity (see methods, Fig. S2A), similar to prior work^1,60,61^. Some remaining ambiguities were resolved using the NeuroPal approach for non-stochastic and unique labeling of neuronal identities^62^. In total, we identified 61 neurons (Fig. S2A-C, Table S3), which, after combining left and right pairs, correspond to 38 neuronal classes. In any single dataset a subset of these were identified, representing ≈ 40% of tracked neurons in freely moving GCaMP datasets, ≈ 38% of tracked neurons in immobilized GCaMP datasets, and ≈ 15% in freely moving GFP datasets (Fig. S2B).

For quantitative measurements of locomotion behavior, we devised an image processing and analysis pipeline that is capable of resolving complex animal shapes to generate multidimensional time-series of behavioral variables, ranging from slow timescale actions to fast timescale granular motor patterns (Fig. S3). Compared to GCaMP signals, traces from GFP datasets acquired under freely-moving conditions revealed only lower variance fluctuations (Fig. S4A-E). Nevertheless, these low variance signals had structure, like correlated noise induced by motion; these recordings thus served as controls for motion artifacts in later analyses.

### Low dimensional and distributed brain activity represents the worm’s major action sequence

We aimed to study neuronal activity underlying baseline exploratory behaviors of freely crawling worms. Thus, we designed an assay with a minimum of external time-varying sensory inputs. Under imaging conditions (n=36 GCaMP, n=8 GFP animals), and for the duration of our recordings, animals exhibited stable forward-crawling, backward-crawling, and turning patterns that enabled them to effectively explore the arena and providing a rich resource of spontaneous behaviors (Fig. S3A-D).

We estimated that 80% of tracked neurons were potentially active because the variance in their activity was above the 95 percentile of the variance of activity of neurons in GFP datasets (Fig. S4C-E), and which exhibited comparable fluorescence levels (Fig. S4F). Moreover, we observed highly synchronized activity patterns among these active neurons (Fig. 1A). This was quantified using Principle Component Analysis (PCA), which captures orthogonal components that explain, in descending order, the most covariance in the data. The top 3 components explained more than 50% of variance (Fig. 1B), indicating low dimensionality of whole-brain activity. Qualitatively, projecting neuronal activity patterns into PCA space revealed a manifold aligned with the action transitions (Fig. 1A, E; Supplemental Video 1).

**Figure 1.**
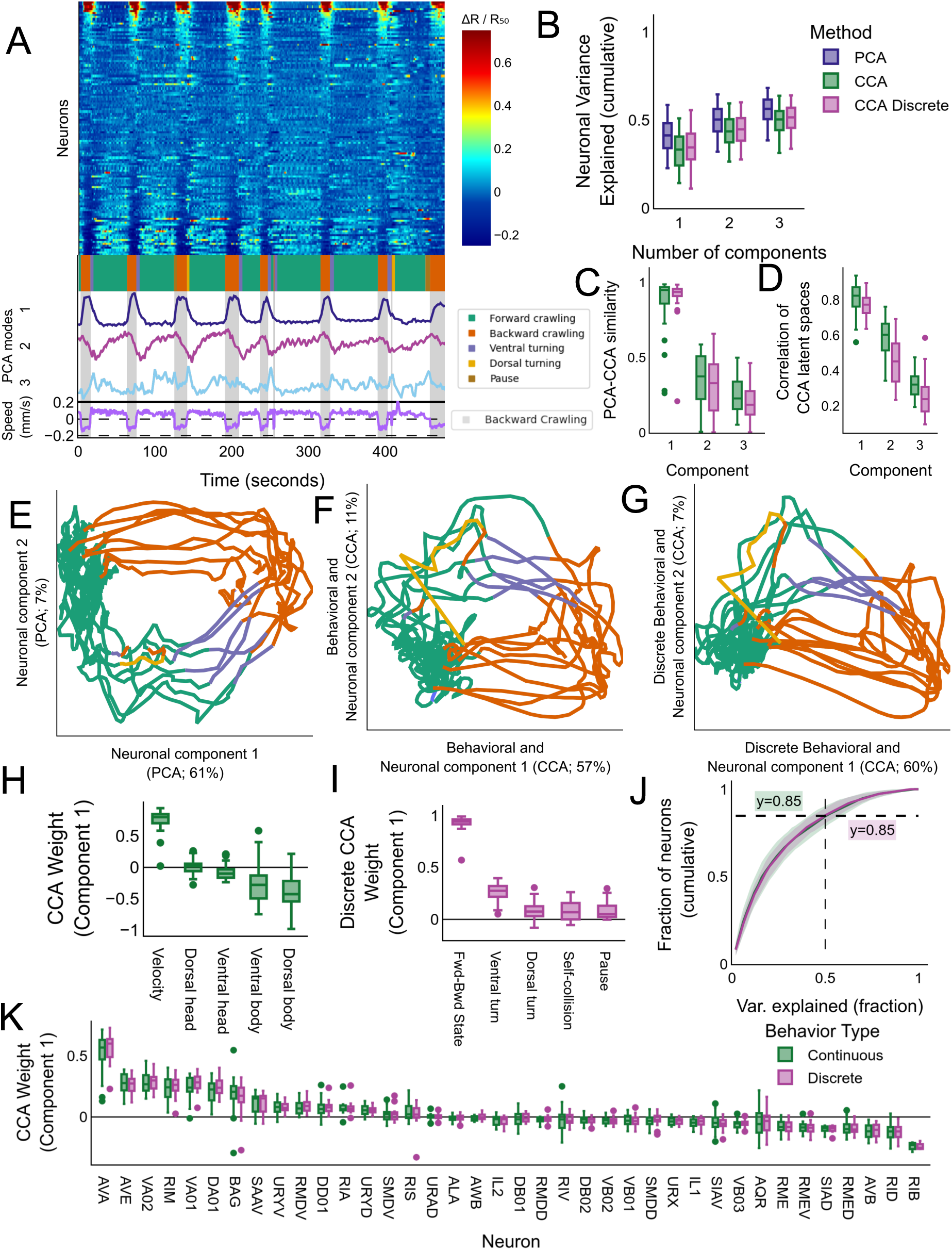
Low dimensional brain activity is dominated by high-level action transitions. **(A)** Heatmap of neuronal activity (ΔR/R50, see colorbar) from an example recording of a freely moving animal showing the activity of 121 tracked neurons across time. The traces are sorted according to the first principle component weights, which reflects the forward-backward crawling transition. Below the heatmap the corresponding trace of the first three principle components and worm velocity (signed by forward and backward crawling) are plotted along with an ethogram indicating the repeated action sequence. **(B)** Variance explained of brain-wide neuronal activity across datasets (n=36) by top components of three different dimensionality reduction methods: neuronal only (PCA), continuous behavioral and neuronal (CCA), and discrete behavioral and neuronal (CCA discrete)._4_ **(C)** Cosine similarity between components of different dimensionality reduction methods across datasets (n=36), comparing individual PCA components to the corresponding component in CCA or CCA discrete. **(D)** Correlation of latent space generated from neuronal and behavioral data, ordered with decreasing correlation. **(E-G)** Examples of neuronal phase space for brain-wide neuronal activity (PCA, **E**), continuous behavioral and neuronal activity (CCA, **F**), and discrete behavioral and neuronal activity (CCA discrete, **G**). **(H-I)** Behaviors used to build top continuous (**H**) CCA or discrete (**I**) CCA component. **(J)** Percent variance of neuronal activity explained by top 2 CCA or CCA discrete components per neuron across datasets (n=36). Traces show mean, shading shows standard deviation. **(K)** The contribution of different identified neurons (left and right pulled) to the top CCA component (weights). Both methods of calculating CCA are shown (n=36 freely moving datasets; see Table S1 for number per neuron). No differences are signficant using a bootstrapped ttest (see Methods). All panels with boxplots show median, interquartile range, and 1.5 times the interquartile range.

To quantify this alignment, we searched for patterns that are shared between behavioral and neuronal data. We therefore performed Canonical Correlation Analysis (CCA)^63^, a method that identifies linear projections with maximal correlations between datasets. Projecting neuronal activity into these low dimensional PCA or CCA spaces revealed similar manifolds, reliably reflecting the forward-backward-turning action sequence (Fig. 1E,F). In particular the top mode of both PCA and neuronal CCA were nearly identical (Fig. 1C). The top two CCA components successfully captured latent dimensions shared in both the neuronal and behavioral data (Figs. 1D, S5A), motivating us to focus on these components. The top CCA behavioral component was composed of instantaneous forward and backward crawling velocity with some contribution from body curvature (Fig. 1H). The second component was composed mainly of body curvature (Fig. S5B). Both components successfully captured the variance of these behaviors (Fig. S5C). Likewise, these components explained a large portion of the variance of the entire neuronal dataset (Fig. 1B). Thus, the low dimensional neuronal manifold, computed either by PCA or CCA, encodes the instantaneous behavior of the animal.

Do low dimensional population dynamics encode continuous behavioral metrics or rather discrete switches between actions? To investigate this question, we performed CCA using discretized action labels. Qualitatively, this analysis showed strikingly similar results to the continuous case (Fig. 1G). A large amount of co-variance in brain activity can be explained by action state alone, and continuous behavioral metrics explain little additional neuronal variance (Fig. 1B). Similar to the continuous CCA method, the top discrete CCA mode is dominated by the forward-backward state switch (Fig. 1I), which is also well reconstructed (Fig. S5D). The second mode has a greater contribution from turning and self-collision, which occurred during high-curvature body postures (Fig. S5B), and successfully reconstructs the ventral turn annotation (Fig. S5D), which were much more common than dorsal turns in our recordings (Fig. S3B). Our results are consistent with the conclusion that a low dimensional manifold largely reflects the progression through the worm’s discrete action states (forward-reverse-turn).

We further found that this manifold is represented in a widely distributed manner at the single-neuron level. Approximately 15% of the neurons recorded having 50% or more of their variance explained by the first two components calculated by either CCA or CCA discrete (Fig. 1J, lines are nearly identical). Further investigating the neuronal constituents of the top CCA components, we found it to be composed of multiple known neurons associated with forward or backward crawling (Fig. 1K), consistent with the activity patterns of these neurons during each action (Fig. 1A). These neurons include the AVA, RIM, and AVE interneurons and DA01 and VA02 motor neurons previously implicated in backward crawling^1,49,64^, as well as AVB, RIB, and RID interneurons and RME motor neurons previously implicated in forward crawling^1,64,65^. The second CCA components are composed of turning neurons; specifically, ventral turns in CCA component 2 and the turn associated neuron RIV^49,66^ (Fig. S5E). Additional known turning neurons like SMDV and SMDD^1,28,49,67^ were also associated with these components, but only to a modest extent (Fig. S5E).

In summary, our analyses uncovered a low dimensional manifold encoding the high-level action sequence of forward-backward-turn behaviors. We further find that this manifold is both dominant and widely distributed across the brain of freely moving animals.

### Low dimensional brain-wide activity is dominated by intrinsic dynamics

What could be the origin of these brain-wide neuronal activity patterns? In freely moving animals, there could be a complex mixture of internally generated activity, external sensory inputs (exafferent sensation), sensation of body posture (proprioception), and other sensory signals caused by the animal’s movements (reafferent sensation). Here, we chose assay conditions largely devoid of time-varying exafferent stimuli (see discussion). In order to distinguish whether neuronal dynamics in freely moving animals are internally driven or derived from proprioception or reafferent sensation, we systematically compared the neuronal activity of freely moving to immobile animals^1,28,60^, which lack these sensory inputs.

We first observed a qualitatively similar dominant pattern of correlated neuronal activity (Fig. 2A,B). This revealed a low-dimensional manifold, as shown previously for immobilized animals^1^ (Fig. S6), although the timing of transitions is different across conditions as also shown previously^1,68^. Of the 61 identifiable neurons (38 neuronal classes after left/right combination), ≈ 71% of neurons that we could identify had a similar relationship to the dominant component (PC1) in both conditions (”Intrinsic” category, Fig. 2C,D). Such a large proportion of neuronal activity that is independent of motor execution is consistent with intrinsically generated dynamics.

**Figure 2.**
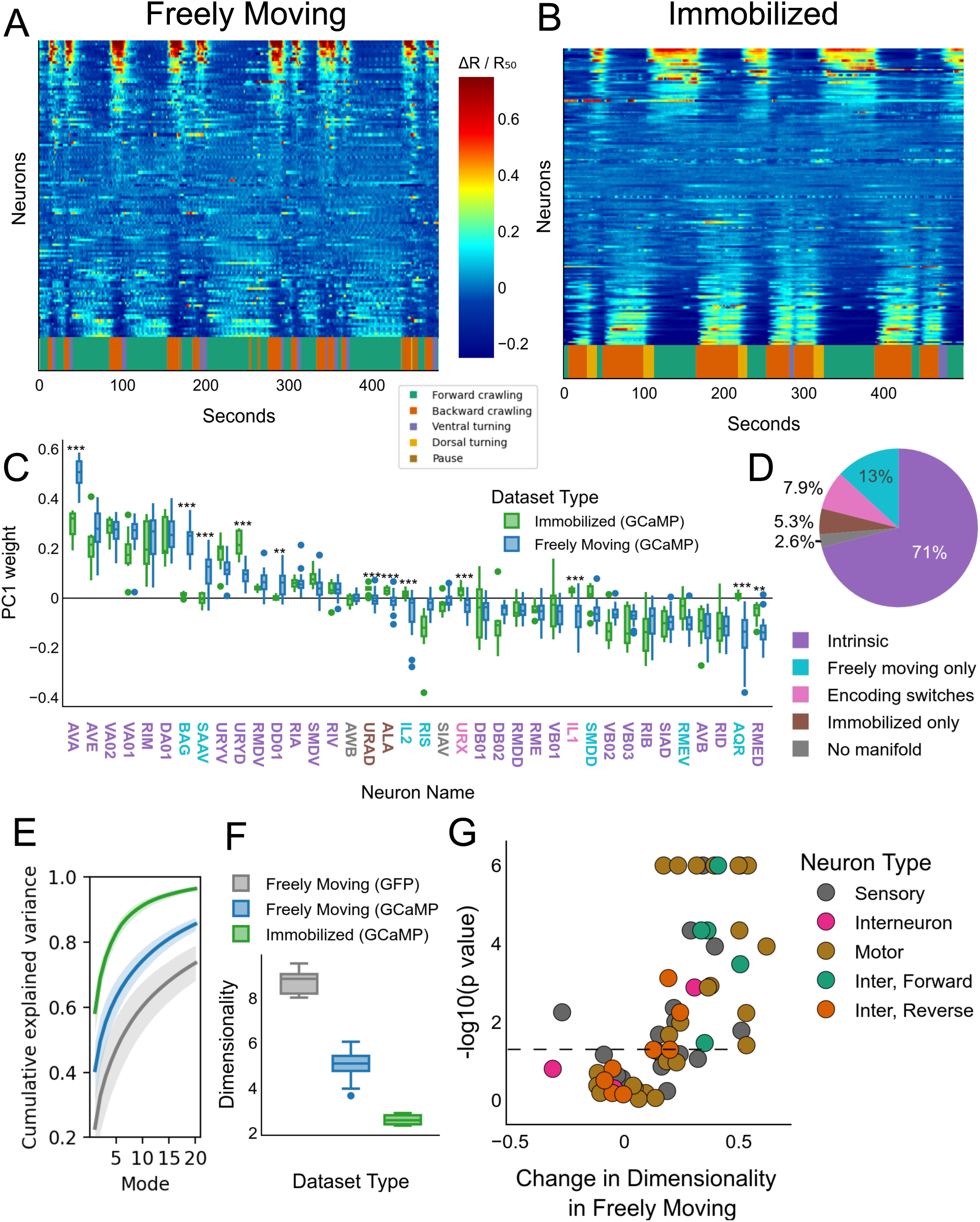
Whole brain activity is dominated by intrinsic activity. **(A-B)** Heatmap of neuronal activity (ΔR/R50, see colorbar) across time of example recordings from two animals, one freely moving (**(A)**; 118 neurons) and one immobilized (**(B)**; 161 neurons). The traces are sorted according to the first principle component weights. The different behavior states are annotated across time (below). For immobilized animals the behavior annotations are based on neuronal activity of motor command neurons (see Methods). **(C)** The contribution of different identified neurons (left and right pooled) to the top neuronal component (PC1 weights) across freely moving (n=36) and immobilized (n=10) individuals. Significance is estimated using a permutation test (See methods), and multiple comparison correction was done by controlling the false discovery rate at alpha=0.05. Significance is shown at the following levels: *, 0.05 < p <= 0.01; **, 0.01 < p <= 0.001, ***, p < 0.001. Exact n numbers for each neuron class are reported in Fig. S2C and Table S1. **(D)** Summary of observed changes of neuronal contribution to top neuron component in panel (C) between conditions. “Intrinsic” means that the PC1 weights did not significantly change between conditions, and “No manifold” means that the PC1 weights in the freely moving case are not significantly different from zero; see Methods for detailed definition. **(E)** Cumulative variance explained by all principle components across freely moving GCaMP (blue; n=36), immobilized GCaMP (green; n=10), and control freely moving GFP (gray; n=8) datasets. Traces show mean, shading shows standard deviation. **(F)** Estimated dimensionality for each dataset type (colors as in (C,E)). Each data point is the median across estimation approaches for one dataset (see Methods and Fig. S4G). **(G)** Volcano plot of estimated change in dimensionality of identified neurons, calculated by an increase or decrease in the variance of neuronal activity explained by the first two principle components (x-axis). Significance (y-axis) was calculated via a permutation t-test (see Methods), and corrected for multiple comparison by controlling the false discovery rate at alpha=0.05. Dashed line indicates p = 0.05. Neurons are color coded by their type and activity pattern. For all panels, boxplots show median and interquartile range, whiskers show 1.5 times the interquartile range.

However, we also found neurons whose encoding changed between these conditions. Approximately 26% of identifiable neurons displayed stronger PC1-correlated activity in one condition (Fig. 2C,D; Table S2). Among the neurons with the strongest significant increase in correlation in the freely moving condition (”Freely moving only” or “Encoding switches” in Fig. 2C,D) to the top neuronal component were gas-sensing neurons BAG, AQR, URX, IL1, and IL2^61,69,70^. We hypothesized that these could be examples of re-afferent perception, which we investigate in a later section. Another neuron class that showed a strong difference is SAAV, which is inactive in the immobilized condition but strongly active during backward crawling. This may be an example of proprioception or another reafferent sensory modality, presumably olfaction^71^. These results agree with a previous study that reported differences in unidentified neurons between freely moving and immobilized conditions^45^. Thus, while many neurons are correlated to the dominant component due to intrinsic activity, some display a similar correlation through reafferent or other sensations.

In summary, we provided evidence supporting the idea that intrinsic dynamics generate the dominant pattern of brain-wide neuronal activity of freely moving animals.

### Higher dimensional activity in freely moving animals

In addition to the intrinsic dynamics, we expected that freely moving animals would also have more complex activity patterns than immobilized animals. Comparing the overall dimensionality of these conditions using several estimation methods (Fig. S4G), we found that datasets from freely moving animals have higher dimensionality (around 4) than that of immobilized animals (around 2; Fig. 2E,F). Guided by this dimensionality estimate in immobilized animals, we defined the top 2 PCA components to be the neuronally-derived manifold. We then checked if this increased dimensionality was present at the single-neuron level. Specifically, we calculated the percent variance explained by the manifold (top two PCA components) in both conditions, and quantified the difference between immobilized and freely moving conditions. Many neurons, including sensory and motor neurons, increased their dimensionality (Fig. 2G). Notably, most backward crawling interneurons did not display a significant increase in dimensionality in freely moving animals. In contrast the forward crawling interneurons display increased dimensionality, which may be evidence for a uniquely complex aspect of the forward crawling command state.

In summary, while most neurons exhibit robust participation in a largely intrinsically generated neuronal manifold (previous section), this result suggests that movement further modulates many of these neurons, increasing the complexity and dimensionality of activity.

### State-dependent encoding of body undulations in multiple motor neurons

In search of interpretable higher dimensional activity, we wondered if these neurons encode more granular motor patterns in freely moving animals. Freely moving animals constantly perform body undulations nested within each of the major action states of forward and backward crawling. Examining the neuronal activity, we found that multiple neurons showed structured oscillatory activity. Specifically, the body motor neurons VB02, DB01, and DD01 displayed oscillations (blue traces, Fig. 3A-C), but others did not, like AVB (Fig. 3D). Importantly, this oscillatory activity did not appear in immobilized animals (Fig. S7A-H). Motor neuron activity is expected to correlate with the undulation of body posture^28,72–75^, however, we observed above that these neurons also display manifold-related related activity (Figs. 2C; 3A-D). To focus on the granular activity, we decomposed the traces of these neurons into the manifold-encoded part (top 2 PCA components), termed the global component, and its residual (See methods). Triggering the global component of these neurons (red; Fig. 3A-D) to forward-backward transition, we found that it is associated with the major action sequence (Fig. S7I-L). Triggering the residual component (Fig. 3E-H, top) to body posture (see methods) revealed that the residual activity patterns of VB02, DB01, and DD01, but not AVB, are strongly associated with neck bends during forward crawling. Moreover, comparing the encoding of VB02 to that of DB01 and DD01, revealed a phase shift according to their innervation patterns of ventral versus dorsal body muscles respectively (Fig. 3E-H, top). All together, we observe neuronal activity that is multiplexed encoding both the major action sequence and instantaneous body posture.

**Figure 3.**
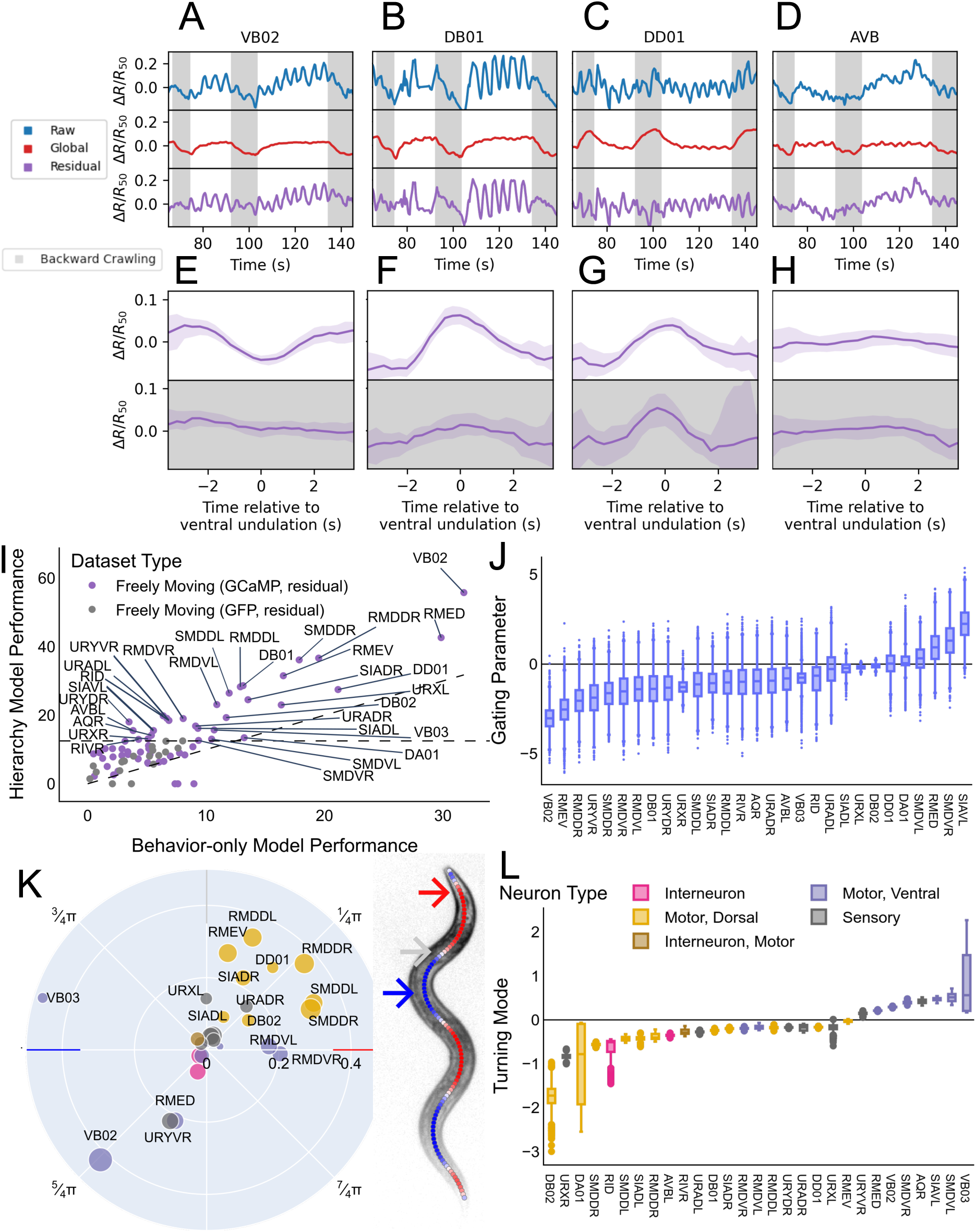
Multiplexed and hierarchically organized neuronal activity. **(A-D)** Example of neuronal traces of VB02 **(A)**, DB01 **(B)**, DD01 **(C)**, and AVB (L/R averaged) **(D)**. Shown are the raw trace (blue) and its decomposition into the global manifold (PC components 1 and 2; red), and the residual activity (purple). **(E-H)** Undulation-triggered averages of the residual activity of the neurons in (A-D), separated by forward- versus backward-crawling states. Shown are VB02 (**(E)**; forward: n=36/1765; backward: n=36/739), DB01 (**(F)**; forward: n=18/926; backward: n=18/358), DD01 (**(G)**; forward: n=19/942; backward: n=19/382), and AVB, left/right averaged (**(H)**; forward: n=35/1727; backward: n=35/727) residual activity. n numbers correspond to number of worms / events. **(I)** Comparison of model performance for behavior-only (x-axis) and hierarchical (y-axis) models. Neurons identified in the control GFP neurons are confined near the origin (gray), which define the significant threshold (dashed horizontal line). Diagonal line indicates equal performance of both models; activity of neurons above this line is better modeled by the hierarchical model. **(J)** Posterior distribution of the gating parameter, the model parameter corresponding to hierarchy (See Methods). Positive values correspond to a hierarchical gating effect by the backward command state, i.e. the residual signal displays behavioral encoding only during the backward command state. In the same way, negative values correspond to gating by the forward command state. **(K)** Best-fit phase relationship between body undulation and neuronal oscillation for all motor neurons that are improved by the hierarchical model. Radial axis indicates strength of oscillation encoding (see Methods). Dot size indicates improvement by hierarchical model as in (I); see also Fig. S7M. Motor neurons are color-coded based on body segments that they innervate. **(L)** Posterior distribution of eigenworm 3, which has been shown to correspond to turning behaviors. Colors are the same as in (K). Boxplots (J, L) show median, interquartile range, and 1.5 times the interquartile range. N numbers for each neuron class are provided in Fig. S2C and Table S1.

We next asked whether the higher level behavior state interacts with the encoding of the body posture. We found that DD01 encodes body bends independent of behavioral action state (Fig. 3G), reflecting the consistent undulation of the body across both forward and backward crawling. In contrast, we found that VB02 and DB01 encode body bends preferentially in the forward action state (Fig. 3E,F). Thus, the activity of VB02 and DB02 is gated by the major action sequence, meaning they encode undulations specifically during forward crawling. Therefore, we hypothesized that the manifold, which encodes an upper hierarchical level behavior (forward and backward crawling), gates neuronal activity related to a lower level behavior (undulation).

### A Bayesian modeling approach reveals brain wide hierarchical encoding of ongoing behaviors

To quantify observations of state-dependent behavioral encoding (Fig. 3E-H) and to test whether hierarchical gating is a more general phenomenon, we took a systematic Bayesian modeling approach. We aimed for a model that can explain hierarchical gating solely based on neuronal activity patterns without external behavioral labels, thus representing a neuronal correlate of a behavioral hierarchy. Further, a useful model should be able to indicate (i) the degree to which individual neurons are gated by the neuronal manifold, (ii) during which brain state gating occurs, and (iii) which behavioral features are encoded by residual activity.

We therefore calculated residual activity for each neuron as in the previous section, and then searched for behavioral correlates in these signals. Given the prominent body undulation encoding we found in DD01, VB02, and DB01, we focused on the eigenworm representation of body posture, where the first two components represent simple body undulation during forward and backward crawling^76^. The third and fourth eigenworms were shown to reflect a turning mode that includes behaviors like post-backward crawling turns^76–78^. We built three Bayesian models: neural traces as a function of behavior alone, or of behavior gated by the global neuronal manifold, and compared the performance of these two models to a third null model which does not use behavior at all. Comparing to a null model, i.e. a flat line model, removes neurons that have no residual activity.

### The manifold gates behavioral encoding of many neurons

To obtain an estimate for the degree of gating, we calculated for each neuron the improvement of each model type above the null model (Fig. 3I): “Hierarchy Model Performance” (y-axis) is the improvement of the hierarchy model and “Behavior-only Model Performance” (x-axis) is the improvement of the Behavior-only model. It was important to carefully control for motion artifacts, which may also be correlated to behavior variables, especially body undulations. We therefore defined the boundaries of non responding neurons by fitting the same model to traces from identifiable neurons from control GFP recordings (Fig. 3I). This analysis statistically confirmed that VB02 and DB01 not only encode body undulations, but are also strongly gated via the global manifold. Specifically, the hierarchical gating model captured the activity for both neurons very well and outperformed the model without the gating component (y axis; Fig. 3I). Many neurons showed strongly hierarchically gated behavior encoding, such as RID, VB02, DB01, RMEV/D, RMDV/D, and SMDDL/R, whereas others such as DD01, SMDVL/R, DA01, VB03, and DB02 are less hierarchically organized (Figs. 3I, S7M). Notably, the strength of gating via the manifold is independent of participation in the manifold i.e., not all gated neurons exhibited an obvious global component. For example, SMDDL/R and RMDDL/R participate only slightly in the manifold (variance explained is ≈ 25%), but they are very strongly hierarchically gated (Fig. S7M). Thus, this analysis uncovered behavioral encodings in multiple neurons which are hierarchically gated to varying degrees.

### The forward manifold state displays the majority of gated behavioral encodings

In our model, hierarchical gating can occur during the forward or backward command state of the manifold. We therefore examined the model’s gating parameter indicating the directionality and strength of gating (see methods). For example, the encoding of oscillations in the model of VB02 and DB01 is strongly active during the forward command state, as expected from their state-specific triggered averages (Fig. 3E,F). Altogether, we observed an enrichment of neurons that are gated during the forward command state (Fig. 3J), e.g. VB02, DB01, RMEV, RMDD, and RID.

### Gated behavioral activity encodes a spectrum of body undulation phases

Next we investigated how the residual signals are coupled to simple body undulations in the hierarchically gated neurons. This revealed stereotyped ventral-dorsal phase relationships (Fig. 3K). The ventral head-curvature encoding VB02 and RMED had opposite phase with respect to the dorsal DB01-2, RMEV, SMDDL/R, and RMDDL/R neurons, as expected from the literature^28,44,79^. This analysis also revealed more subtle phase shifts between the encoding of the neurons. The VB03 neurons had an intermediate encoding, between the dorsal DB01 and ventral VB02 neurons, consistent with its innervation of muscles further down the body^80^. Examining the motor neurons as a whole, we found a sequence of gradually changing phase encoding: Starting from the RMDVL/R neuron pair, followed by the SMDDL/R, leading to DB02 and DB01, followed by SIADL/R and RMEV. Notably the RMDV undulation phase encoding was not ventral or opposite to that of RMDD, as expected based on muscle innervation pattern, but instead was more similar to phase encoding of dorsal neurons. In total, we discovered that the residual activity of multiple neurons encodes detailed information about a wide range of phases of the body undulation.

Finally, we investigated the encoding of turning behaviors. Consistently, dorsal or ventral muscle innervating neurons encoded the appropriate sign of the ventral (Fig. 3L) and dorsal turning modes (S7O). Some neurons like SIAVL, SMDVL/R, RID, and AVBL show substantial activity correlated with turning behaviors, but very little simple undulation correlated activity (Figs. 3K,L, S7O). Other neurons, like the ventral-correlated VB03 and VB02 and the dorsal-correlated DB01, DB02, SMDDL/R, and RMDDL/R, show both undulation and turning-related activity (Figs. 3K,L, S7O).

In total, this analysis suggests that the manifold hierarchically gates the behavior-related activity of many neuron classes. This potentially allows the animal to execute granular motor patterns (body undulations and turns) at the proper time within high-level action states (forward and backward crawling states).

### Re-afferent perception is apparent in low dimensional global activity

In the previous section, we proposed a function for the global neuronal manifold. However, not all activity correlated with this component needs to have the same function or in fact be intrinsically generated. For example, we found the activity of multiple gas sensing neurons to follow the global component, but only in freely moving animals (Fig. 2C,D). Behavior-correlated activity could stem from re-afferent sensation of fluctuations in local gas concentrations, such as of self-produced CO_2_ and respired O_2_, as previously found for the BAG neurons^40^. To test this hypothesis, we first performed whole-brain recordings of freely moving animals, then immobilized the same individuals, and recorded their neuronal responses to external O_2_ stimuli (See methods). These experiments revealed a large number of O_2_ sensitive neurons that also react to either forward or backward crawling movements under freely moving conditions (Fig. S8A-F). We first replicated prior work^40^, showing that BAG ramps its activity during backward crawling in freely moving animals (Fig. 4A) but is not correlated with the backward crawling command state in immobilized animals (Fig. 2C).

**Figure 4.**
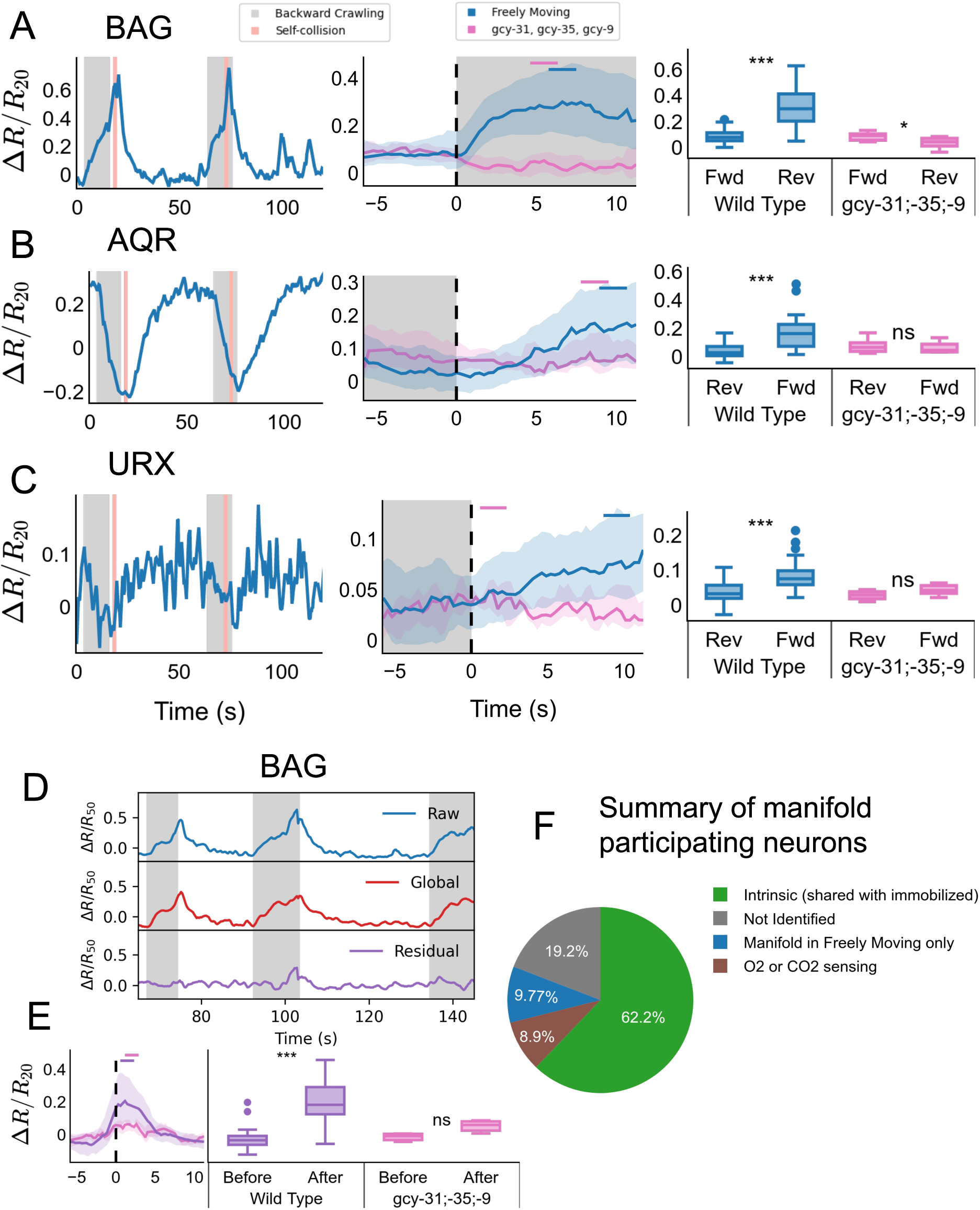
Reafferent sensory information contributes to the global manifold. **(A-C)** Left column: Example raw traces for BAG **(A)**, AQR **(B)**, and URX **(C)**. Middle column: triggered averages for each neuron class across wild type and O_2_/CO_2_ sensory mutant (gcy-31; −35; -9) datasets. See Table S1 for number of ids and events per neuron and per strain. Events within one dataset were averaged, and line shows mean and shading shows standard deviation across datasets. All analyses pool left and right neuron pairs. Small bar refers to the time window used for ttest (see Methods). Significance is shown at the following levels: *, 0.05 < p <= 0.01; **, 0.01 < p <= 0.001, ***, p < 0.001. Right column: comparison of the mean of preand post-backward crawling activity in each genotype. Boxplots show median, interquartile range, and 1.5 times the interquartile range. **(D)** Example trace of the O_2_/CO_2_ sensing BAG neuron. Below are the global component (PC modes 1 and 2; red) that responds to reversal behavior state and the residual component component (purple) that responds to self-collisions. **(E)** The average activity of the BAG neurons in response to self-collision. Shaded area represents standard deviation. **(F)** Summary of classification of manifold-participating neurons in freely moving datasets (n=36). Manifold participation is defined as have >50% of the variance of the neuronal activity explained by the manifold (first 2 PCA modes). Not identified neurons might correspond to one of the other categories.

Further, we found that O_2_ upshift sensing neurons like AQR and URX^69^ (Figs. 4B,C; S8B,F), as well as AUA^81^, IL1, and IL2^61^ (Fig. S8C-E) are activated by transitioning to forward movement. We further confirmed that the freely moving activity of these neurons is sensory derived using a mutant strain defective of the known O_2_ and CO_2_ sensory pathways (gcy-31;gcy-35;gcy-9^69,70^; Fig. S8A-F). In this genetic background the responses of the O_2_ upshift neurons were abolished (Figs. 4B,C, S8B-F, pink). Moreover, we found the activity of BAG neurons to spike when the worm’s nose collides with the body, which was highlighted by the global-residual decomposition discussed in the previous section (Fig. 4D). This response was consistent across individuals, and also abolished in the mutant background (Fig. 4E). Altogether, these results show that the re-afferent gas signal is broadly encoded in multiple O_2_ and CO_2_ sensing neuron classes (Fig. 4F).

We next wondered if the global component also gates the activity of these gas sensory neurons, as we showed previously for BAG neurons that get suppressed via corollary discharge from the backward active RIM neurons^40^. In wild type animals, there is a complex mixture of sensory inputs and internally driven dynamics, such as motor commands and corollary discharge. Gas sensory mutant animals, which lack sensory activity, allow these effects to be disentangled: in these mutants, the BAG activity was unmasked from sensory driven activity and revealed the inhibitory corollary discharge signature (right column, Fig. 4A). However, we did not detect a similar signature of corollary discharge during forward crawling in other neurons.

In summary, about 60% of the neuron classes active in the global manifold can be identified as shared with the immobilized animal, and a further 9% are active due to O_2_ or CO_2_ re-afferent perception (Fig. 4F). We show that even under these conditions, the animals’ brains engage in embodied perception, being sensitive to the sensory consequences of their own movements.

## DISCUSSION

In this work, we recorded brain-wide neuronal activity in freely moving animals, and discovered low-dimensional structure across the activity of many sensory-, inter-, and motor neurons. This structure can be modeled as a neuronal manifold, which faithfully reflects the animal’s cyclical progression through an action sequence of motor programs. Our findings support the idea that this signal is largely intrinsically generated, with a minor contribution from re-afferent sensory activity. We further find residual neuronal activity in many neuron classes which are hierarchically nested or gated via the low dimensional manifold and which encode granular posture dynamics. Thus, we found hierarchically organized neuronal activity corresponding to a hierarchy of motor behaviors. This work shows a potential hierarchical organizing principle: information about the current action state is broadcast across the brain via a low dimensional manifold to orchestrate fine-scale activity in the motor periphery.

### Intrinsic neuronal dynamics dominate brain activity

We observe rich neuronal dynamics encompassing a large fraction (≈ 80%) of all tracked neurons (Fig. S4E). A striking feature of this neuronal population dynamics are correlations among many sensory-, interand motorneuron classes. For most neuronal cell types, this correlational structure, quantified by PC weights, was reproducible in immobilized conditions (Fig. 2C). Moreover, both conditions showed frequent transitions in these correlated activity patterns. Importantly, in the immobilized condition animals are not only deprived of external acute stimuli, but also lack any time-varying re-afferent inputs or proprioception of their own movements. This result supports the idea that these dynamics are intrinsically driven. This does not exclude that the environment can influence these dynamics and their associated behaviors. For example, it was shown that constant or only slowly-changing arousing factors such as time-off-food^49^, hunger^82^, or experimental conditions such as excitation light^68^ can affect the frequency of behavioral or brain state transitions. However, such factors cannot explain exact transition timing and the observed correlational structure in neuronal dynamics.

The origins of transition timings are unknown, but intrinsic stochastic mechanisms have been discussed^50^. The neuronal connectome can provide some insights into the potential mechanisms underlying the correlational structure: we previously reported that the rich-club architecture of the connectome supports this structure^60^. A set of highly interconnected rich-club neurons was crucial to establish brain-wide activity correlations. In the present study, consistently, the rich club interneurons AVA, AVE RIM, AVB and RIB were found active and were among the major contributors of low dimensional dynamics (Figs. 1K, 2C, S5E).

Taken together these lines of evidence strongly suggest that these dynamics are most likely established by intrinsic mechanisms.

### Low dimensional population dynamics represent the major action sequence of *C. elegans*

Based on these strong and stable correlations, it is possible to model brain dynamics as a low dimensional manifold, which we demonstrate here for the first time in freely moving worms. We show that this manifold is strongly correlated with the major behavioral action sequence of the worm tracing repeated cycles of forward crawling, backward crawling, and turning (Figs. 1, S6, Supplemental Video 1). Indeed, a large body of work showed that most neurons we found contributing to the manifold are functionally implicated in these action states^60^ (reviewed in^83^; see also references collected in Table S1 in^1^). More recent work in *C. elegans* has demonstrated mappings between various behavioral metrics and neuronal activities, revealing neuron class specific behavioral tuning curves^1,44–46,65^. However, the behavioral mapping of the manifold itself remained unclear. Here we use Canonical Correlation Analysis (CCA) to establish a linear mapping between low dimensional neuronal population activity and a set of such behavioral metrics. We show that both neuronal activity and behaviors are well modeled by the combined neuronal-behavioral manifold obtained via CCA (Figs. 1B,D, S5C,D). Surprisingly, using discrete behavioral metrics e.g., forward-backward state, performed nearly equally well at explaining neuronal activity. This result could be partially explained by the bimodal distribution of crawling velocity resembling the binary Forward-Backward crawling state; both metrics contributed most to the first CCA mode in the two approaches (Fig. 1H,I). Alternatively, we propose that a discrete action state could be the prime information that is broadcast across the brain, providing a context for individual neuron types to execute their specific functions.

### State-dependent gating of neuronal activity by low dimensional population modes establishes behavioral hierarchies

While individual neuron classes likely all execute specific functions, their functional activity appears to be coupled to the global population modes. Our Bayesian model provides a quantitative basis for these observations: the activity of many motor neurons exhibits residual activity patterns not explained by the low dimensional manifold. However, the behavioral mappings of these residual activities were modeled best upon hierarchically nesting into the manifold structure (Fig. 3I), which conveys action state information. Notably, commonly such mappings between behavior and neuronal activity rely on manually defined behavioral state labels (forwardbackward-turning) in order to obtain neuron specific behavioral tuning curves^1,44–46,65^. In our hierarchical model, since we used PCA modes, this state dependency is instead derived from low dimensional neuronal population modes, not requiring external behavior labels. Therefore, non-linear state dependencies in behavioral encodings can be explained directly by intrinsic brain activity, suggesting a neuronal top-down hierarchy corresponding to a behavioral motor hierarchy.

Hierarchical nesting is very intuitively demonstrated with forward crawling B-class motor neurons (B-MNs) as examples. These motor neuron types are inactive during backward crawling and oscillate during the forward command state (Fig. 3A,B,E,F), to establish the undulatory crawling gait that propels the animal in forward direction. Reconciling our data with the available literature on these neurons, we can propose a detailed mechanistic model of hierarchical gating. The rich club neuron AVB is a major contributor to the low dimensional manifold (Fig. 1K, 2C). It represents a bottleneck in the connectome between the brain and the pool of all B-MNs^80,84^ and conveys a descending motor command to them^60,64,85^. The proprioceptive properties of B-MNs^72,73^ likely together with central pattern generator (CPG) circuit elements^28,85^, then maintain oscillations in coordination with other body segments. Our finding that the B-MNs DB01 and VB02 do not oscillate but retain their global signal component in immobilized worms further supports this model: in immobilized worms, the effect of the descending forward motor command is detectable, while in crawling animals this signal is modulated, according to this model, by CPGs and proprioception.

We show that hierarchical gating of higher dimensional residual activity patterns is not unique to B-MNs but extends to many other neuronal classes, in particular head motor neurons (Fig. 3I-L). These exhibit more complex connectivity patterns with the rich club^80,84,86^. Two recent studies focusing each on different subsets of head motor neurons, provide some mechanistic insights how hierarchical gating could be achieved in these cases^28,31^. Interestingly, many neurons exhibit more complex activity patterns in freely moving worms, including motorneurons and particularly interneurons associated with forward crawling (Fig. 2G). This is consistent with a recent whole brain imaging study reporting a diverse continuum of behavioral representations in the *C. elegans* brain^44^.

Our hierarchical modeling approach reveals an interpretable organizing principle that demonstrates how such diverse activity patterns are orchestrated to achieve robust and temporally organized behaviors. We propose a model, where rich club interneurons broadcast behavioral state information and/or other behavioral metrics like crawling velocity across the brain. This provides structured context for orchestrating neuronal activity patterns at the periphery to execute neuron specific functions. It is important to consider that each neuron class participating in these low- and high-dimensional neuronal dynamics have unique connectivity patterns with the rich club; this potentially enables specific communication channels (termed subspaces). For example, B-MN’s synaptic inputs are largely dominated by AVB; thus AVB to B-MN connectivity represents the major communication subspace to this neuron class. Conversely, head motor neurons with more complex connectivty patterns^80,84,86^ likely communicate with the rest of the brain via more diverse subspaces. Moreover, our recent work showed that dynamical neuronal interactions across the *C. elegans* connectome are recurrent and indirect^60^. A single global manifold, obtained via PCA or CCA, should therefore be seen as a computationally convenient approximation of potentially more diverse neuronal communication subspaces, and each neuron class likely receives a different variant of the global signal. We here chose these dimensionality reduction methods for their robustness and ease of interpretation.

However, there are many nonlinear extensions of these algorithms, and future studies could apply them to achieve improved manifold representations^87,88^. More broadly, dimensionality may not be the key dividing factor between a manifold and other residual activity. For example, a decomposition could highlight slow dynamics, like Slow Feature Analysis^89^, or separate multitimescale phenomenon in general^90^. Especially when the low dimensional manifold represents slower and higher level actions, as in this study, all of these decompositions may give similar results. A promising area of future work is to test such decompositions to determine if there are interactions between low dimensional subspaces and behavioral encoding.

### Interdependence of low dimensional and residual population activity

Based on theoretical work it could be debated whether the dimensionality of brain-wide neuronal activity is bounded or not (compare refs.^91,92^ with refs.^93,94^); conceptually, the unbounded case comes with the consequence that any discrete cutoff to define low-dimensional manifolds might be arbitrary. We argue here that the neuronal hierarchies we propose operate irrespective of discrete boundaries between low- and high dimensional population activity, although such cutoffs are required in practice for statistical analyses. Here, we chose a dimensionality cutoff of D=2, based on the general agreement of various estimation methods in immobilized recordings (Fig. S4G), which should reflect the best baseline for intrinsically generated low dimensional dynamics.

Coordinated activity across populations of neurons, bounded or not, potentially influences the residual activity of many neurons. These interactions can be even reciprocal: head motor neurons have been shown to also influence behavioral state and brain state transition frequencies^28,31,66^. Thus, the hierarchies we propose here operate not strictly in a top-down only manner.

### Implications for sensory processing

We found that reafferent sensation of locomotion-induced local CO_2_ and O_2_ fluctuations has a small but significant contribution to behavior-correlated neural dynamics (Figs. 4, S8). Specifically, we found O_2_-sensitive neurons to be activated by forward crawling, and CO_2_-sensitive neurons to be activated by backward crawling or turning, in case of self-collisions. These signals depend on CO_2_ and O_2_ sensory pathways, indicating that they reflect self-sensation. We have previously shown that animals sense their own remnant CO_2_ plume when crawling backward into their previously occupied path^40^. Activation of O_2_ sensory neurons is consistent with an analogous interpretation that worms sense a rise in O_2_ upon crawling out of a O_2_ sink when switching from backward to forward crawling. These reafferent inputs are likely accentuated in the coverglass sandwich configuration used in this study; we recently discussed that this experimental condition likely mimics ethological relevant aspects of aqueous natural environments (for more details see ref.^40^).

Reafferent sensation of metabolic gases might not be the only sensory cues that are strictly coupled to locomotion. Mechanosensory neurons detecting texture of the environment or additional modalities altered by self-made stimuli might behave similarly. In the current study, we chose an isotropic arena to study spontaneous behavioral transitions, i.e. that are not acutely evoked by spatial or temporal fluctuations in the environment. This setting enabled us to focus on the neuronal correlates of such intrinsic behaviors and sole reafferent inputs thereby establishing a baseline for future studies on how animals integrate internal behavioral representations with exafferent sensory experiences. Our study highlights that movement related signals in sensory brain areas should be critically investigated before concluding that these signals are caused by efference copies (see also ref.^7^ for an example in macaques). Notably, *C. elegans* navigating sensory landscapes generate the time-varying sensory inputs crucial for behavioral control with their own movements i.e., by crawling up- and down- odorant gradients and undulating with their nose^48,95,96^. Based on this we expect that exaffernet sensory representations in freely moving worms must be also tightly coupled to locomotion behaviors. Therefore, it is crucial to first thoroughly characterize neuronal activity in isotropic environments so that exafferent and reafferent sensory stimuli can be unambiguously distinguished. Future whole brain imaging studies of animals navigating controlled sensory landscapes should take into account these considerations.

Previous work showed that behavioral state is crucial for context dependent sensory processing^40,97–99^, for example via broadcasting the current behavioral state as corollary discharge or efference copies^35^. Previously, we showed that reafferent CO_2_ stimuli are largely suppressed by backward crawling-coupled tyraminergic corollary discharge signal sent from the rich club neuron RIM to BAG sensory neurons^40^. Thus, the signal in BAG measured in this study is a remnant of incomplete suppression. Moreover, we find a signature of suppression via corollary discharge in the absence of CO_2_ and O_2_ sensory pathways, but only in the BAG neuron class. Disambiguation of exafferent and reafferent signals may occur downstream of sensory neurons by combining correlated information from sensory pathways and the intrinsic manifold. We therefore propose that that the manifold itself can act as a corollary discharge signal in addition to the above mentioned hierarchical motor control.

### Hierarchical gating as a general principle of brain activity organization

We propose hierarchical gating as a more general working hypothesis for investigating the role of brain wide neuronal dynamics in larger model species. In fruit flies, brain wide population modes correlate with salient behavioral states such as running and flailing^2–4^, while a large amount of residual activity across the brain remains to be functionally characterized^3^. In rodent cerebellum, during spontaneous behaviors, low dimensional manifolds of granule cell (GrCs) population activity represent behavioral state, such as active locomotion vs quiet wakefulness, while faster time-scale whisking movements are represented in a distributed fashion across higher dimensions of GrCs activity^100^. In rodent cortex, neuronal manifolds can decode action identity^101^, and low dimensional modes can decode coarse behavioral parameters such as running speed or general whisking activity, while more granular behaviors such as diverse orofacial movements, are represented in high-dimensional population activity^91^. In primate motor cortical areas, low dimensional population dynamics encode aspects of reaching movements^10,102^, while the more granular grasping movements of hands are represented by more complex higher-dimensional dynamics^22^. Taken together, while low-dimensional dynamics generally appear to represent aspects of coarse movements or behavioral states, it is less known how the underlying circuits orchestrate associated granular and faster time-scale behaviors. In these larger animal models, it has not been addressed yet whether brain-wide activity could be organized on a manifold, whether manifold structures connect multiple actions into behavioral sequences, and how faster time-scale behaviors are thereby orchestrated. Here we provide a conceptual and methodological framework that leverages large-scale, simultaneous neuronal activity and behavioral recordings to address these open questions.

## METHODS

### Image acquisition

Worms used in this study expressed a pan-neuronally and nuclear-localized GCaMP7b (or GFP for control experiments) along with a red nuclear-localized mScarlet as a reference marker. Constructs were codon optimized for *C. elegans* expression. The recorded worms were well fed young adult hermaphrodites (first adulthood day, with 0-2 eggs). To prevent bacterial contamination of the pad, worms were first transferred to an unseeded plate to let them crawl for a few seconds out of any co-transferred food remnants and then transferred onto the final assay pad. For the imaging arena, we devised a sandwich configuration where animals are placed on an agarose pad (40.5mm x 36mm x 2.5mm) under a coverglass (ClariTex #1 thickness, 64×50mm). An additional identical coverglass was placed on the opposite side of the agar pad. This configuration stabilized the worms vertically (z-position) and allowed water-immersion optics. The pads were created using 2 % agarose mixed with NGM buffer poured into 3D-printed templates between glass plates, ensuring consistent pad size and flat surfaces. A spacer (2.6mm in height) surrounded the pad and thereby slightly lifted the coverglass above the worm, reducing the pressure on them. This addition, facilitates better spacing between the neurons and improves segmentation performance by the computational pipeline (Fig. S1).

The microscopy system consists of a Zeiss Observer.Z1 inverted microscopy body. Volumetric fluorescence imaging was performed from below through the coverglass, while behavior was recorded through the agar pad. As Objective a Zeiss LD LCI Plan-Apochromat 40x/1,2 Imm Korr with water immersion was used. On the left side port of the microscopy body a Yokogawa CSU-X1 spinning disc was mounted, with a Di01-T405/488/561 dichroic installed. Between the spinning disc and the microscopy body a quadrant-photomultiplier (Applied Scientific Instrument (ASI) PhotoTrack system) was placed. A dichroic short pass mirror @ 620nm was used to direct light beyond 620nm to the quadrant photo multiplier of the PhotoTrack system. The PhotoTrack system was connected to an ASI MS-2000 Controller, which controls the ASI PZ-2000FT XY stage of the microscopy system. In addition a Physics Instruments (PI) P-736 PInano Z-microscopy scanner was used as top plate on the stage to scan the specimen. The PI P-736 scanner was controlled by a PI E-709 piezo controller. On the camera port of the CSU-X1 system a custom beam splitter with 0.5x magnification relay lens system for green and red imaging channels was used. This configuration serves as optical binning, increasing intensity per pixel and reducing the recording area on the sensor, thereby increasing acquisition speed. The beam splitter was built using Thorlabs components. For the beam splitter cube a Semrock FF580-FDi01 dichroic beam splitter and 525/50 and 617/73 band pass filter were used. The images were recorded using two pco.edge 4.2 cameras with Camera Link HS connection. The resulting xy resolution on these cameras was 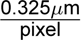 and a region of 900×650 pixels was imaged. As light source for the CSU-X1 fiber port a Oxxius L6Cc combiner equipped with LBX-405nm-180-CSB, LBX-488nm-100-CSB and OBIS-561nm-100 lasers was used. On top of the microscope a machine vision System (Moritex telezentric objective type MML2-HR65D) with 2x magnification, an 850nm coaxial illumination and a Basler ace acA1300-200um camera were installed. For synchronization of all components a Teensy 3.6 microcontroller was used. To control the system a custom Micromanager Plugin was written. All devices where triggered or controlled by the micro controller, which gives synchronous signals to the two pco.edge cameras and the Basler behavior camera, analog outputs for P-736 scanner control and trigger to the MS-2000 stage to trigger position readout. During each z-section of fluorescence imaging, a behavior frame was captured synchronously.

Animals were centered at the head-region under high-resolution optics for fluorescence imaging via the quadrant-photomultiplier controlling the motorized x-y stage^1,51^; volumetric imaging was achieved with the piezo stage scanning in the z-dimension. Volumes were acquired with 24 z slices per volume. Scanning was done using a z step of 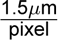, with a flyback at the end of each z-stack. Due to the flyback and subsequent oscillations during the settling of the piezo, the first two (of 24) z slices after each flyback were discarded.

Neuronal activity and behavior were recorded at a volume rate of approx. 3.5Hz and a framerate of 83.3Hz, using an exposure time of 12ms for both modalities. All recordings in all conditions were 8 minutes long, except for seven 4-minute long recordings and one 7-minute long recording in the freely moving GCaMP condition. In addition immobilized gas response recordings were 20 minutes long. These were shortened due to complete tracking failure or tests to decrease bleaching (not shown).

### Immobilized Chip recording

Whole-brain recordings of immobilized worms in an oxygenated microfluidic channel were performed as previously described^1,60^. Young adult worms (0–6 eggs) were immobilized in liquid NGM (nematode growth medium: NaCl (0.3 g/mL), Peptone (0.25 g/mL), Cholesterol (0.5 mg/mL), MgSO_4_ (1 mM), CaCl_2_ (1 mM), and KPO_4_ (2.5 mM)), supplemented with 1 mM tetramisole to paralyze the muscles. Worms were placed in a two-layer PDMS microfluidic device to control O_2_ levels^69^ and align their left-right body axis with the z-scanning dimension using a curved inlet channel (see^41,60,69^ for more details on the chip design). This configuration enabled simultaneous imaging of all neurons in the head and tail ganglia, the retrovesicular ganglion, and multiple motor neurons along the ventral cord. To keep the set of visible neurons consistent between the freely moving and immobilized conditions, neurons in the tail were excluded from the analysis.

The microfluidic device was connected using Tygon tubing (0.02 in ID, 0.06 in OD; Norton) and 23G Luer-stub adapters (Intramedic). A constant gas flow of 21% O_2_ and 79% N_2_ (50 mL/min) was maintained using a gas mixer with mass flow controllers (Vögtling Instruments), controlled via custom-written Micro-Manager software. For general immobilized condition, worms were recorded for 8 minutes and exposed to a constant 21% O_2_ (balanced with 89% N_2_). For immobilized recording with gas responses, worms were recorded for 20 minutes and exposed to the following gas protocol: the worms were habituated to 11% O_2_ (balanced with 89% N_2_) for 4 minutes, then underwent four cycles consisting of 30 seconds in 21% O_2_ and 60 seconds in 11% O_2_. Next, they were exposed to 21% O_2_ for 4 minutes, followed by four cycles of 30 seconds in 11% O_2_ and 60 seconds in 21% O_2_.

For validation of the identities of the gas-responding neurons, individual worms were first recorded in freely moving for 4 minutes and then the same individuals were immobilized with tetramisole (1mM) and transferred to the microfluidic chip for additional recording for 8 minutes. The immobilized worms were exposed to 4 minutes of 11% O_2_, followed by four cycles of exposure of 30 seconds to 21% O_2_ and 60 seconds to 11% O_2_. This immobilized condition exposed the identity of the gas responding neurons (BAG, AQR, URX, and AUA). These identities were then matched between the immobilized and freely moving recordings of the same individual. This method helped define their typical position, shape, and activity pattern in freely moving worms, making it easier to identify them in other freely moving animals.

All microscope and imaging settings matched those used for freely moving experiments, except for longer exposure times of 20 ms and reduced blue laser power of 100µW. The volume acquisition rate was approximately 2 Hz. The reduced light intensity lowered quiescence, which was likely caused by the combination of high light intensity and quiescence induced by microfluidic immobilization^103^.

We initially observed reduced neuronal responses to oxygen shifts in the IL1 and IL2 neurons and suspected the cause was the immobilization agent tetramisole. To address this issue, we immobilized the animals using an alternative method. We used animals expressing the histamine-gated chloride channel HisCl^104^ under the pan-muscular promoter *Pmlc-2*^105^ to inhibit muscle activity. The animals were placed in bacteria-seeded NGM plates with 20 mM histamine for 30 minutes. Histamine (20mM) was also included in the NGM buffer used in the microfluidic channel. This protocol was used for O_2_ stimulation experiments in immobilized animals validating oxygen responses of IL1 and IL2 neurons, shown in (Fig. S8 D, E).

### Behavioral quantifications

To find neuronal correlates between neuronal activity and behavior we devised a pipeline to extract a series of behavioral variables from our behavioral recordings. Both continuous and discretized behavioral metrics were used throughout the paper. Most of these time series were derived from the full body curvature annotation, the kymograph (Fig. S3D). Producing kymographs required a series of image processing steps to generate a head-to-tail centerline from raw images.

### Extracting Centerlines and Kymographs

We binarized our infrared images to obtain a mask of the worm, using a U-Net^106^. For the body postures where the worm is colliding with itself (Fig. S3B) we separately trained another U-Net to be able to extract centerlines. We trained DeepLabCut^107^ to annotate the nose and the tail positions on the infrared images, which served as anchor points to fit a 100 knot uniform spline on the mask using the scipy library in Python^108^. The centerline coordinates of the worm were used to calculate the local body curvature using the following formula^109^ 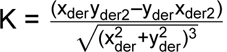

The resulting timeseries of the 100 knots curvature values across the recordings were finally calibrated for dorsal and ventral bending, by identifying the vulva to determine the ventral side of the body. The result is a kymogram (Fig. S3D) which gives detailed insights into the worm’s granular motor patterns. Previous work showed that local curvature directly relates to muscle activity^110^. Qualitatively, the animals’ behaviors did not differ compared to the ones observed under conventional laboratory conditions, with the exceptions of a subset of backward crawling actions, during which the animals curled-up their tail, probably caused by the higher agarose concentration and the agarpad-coverglass sandwich condition used in this study.

### Hilbert transform

In order to study curvature related neuronal activity across individuals, we designed a procedure to calculate triggered averages for this continuous variable, the instantaneous phase of each body bend. Specifically, we used the Hilbert transform^111^ to decompose the kymograph of body curvature into a carrier wave, instantaneous frequency, and instantaneous phase. This procedure is qualitatively similar to simply fitting a sine wave to the raw curvature time series and extracting the phase, but the Hilbert transform also accounts for time varying amplitude and frequency. “Onset” points were then defined as every time this phase returns to the same point. The phase continuously increases from 0 to 2*π* during forward motion, and decreases during reversals, and thus onsets were defined to occur every time the phase passed a threshold of *π*.

### Continuous Behavior Annotations

The velocity of the worm was extracted from the absolute tracking stage position (Fig. S3A) Velocity is calculated by first taking the stage position at the same frequency as the volumetric imaging, aligned with the flyback. The main calculation to take the derivative using numpy’s gradient function^112^. Then, outliers are removed by taking the rolling mean (window of 10 frames) and removing values more than 2 standard deviations away from this smoothed time series. Finally, any remaining large values are clipped to 0.5 mm/s. Its distribution was quantified to verify that under whole brain freely moving conditions our worms have normal locomotion.

Eigenworms (EWs) were calculated by performing PCA on the kymographs^76^. We excluding 10 segments at the head since complex head movement captured by our high-resolution method typically distorted EWs compared to previous work^76,77^. We also removed 10 segments from the tail; these segments were typically very noisy due to the progressive lack of contrast towards the tail tip, which tapers and becomes very transparent. For the cross-individual Bayesian model in Figs. 3I-L and S7M-O, the PCA decomposition was standardized across individuals by concatenating their kymographs.

Further continuous annotations were generated and used in Fig. 1. Ventral or Dorsal head curvature were defined as the mean of body segments 2-9, but only taking positive or negative values, respectively. Ventral or Dorsal body curvature was defined in the same way, as an average of segments 10-90 of only the positive or negative curvature values.

### Discrete Behavior Annotations

A very well described motor action sequence of the *C. elegans* worm is the Forward-Reverse-Turn transition. We automatically annotated reversals from the kymograms using the phase relationship between eigenworm 1 and eigenworm 2 projections, which form a quadrature pair^76^. Forward and backward crawling states correspond to the sign of the angular velocity which was obtained from the cross product of eigenworm 1 and eigenworm 2. From this we quantified the duration of the forward and backward states (Fig. S3C).

Further discrete annotations were generated and used in Fig. 1. The pause state was defined based on a speed threshold of 0.01 mm/s; very short pauses of less than 0.5 seconds were then filtered out.

Worms crawl on one of their sides (left or right) and therefore their turns can be either in ventral or dorsal direction. To be able to distinguish those, we manually labeled the ventral side of each worm by vulva position and checked that the worms did not flip sides throughout the recording. After this, we signed ventral curvature as positive, and dorsal curvature as negative. The signed curvature (ventral – dorsal) was used to identify turn events and label them as ventral or dorsal, with the aim to find neuronal candidates that would correlate to dorsal or ventral bends, as shown previously for SMDD and SMDV head motor neurons respectively^1,28^. Turns were defined as starting at the end of a reversal and continuing until the nose bend curvature changed sign. Specifically, the nose curvature was defined as the mean of body segments 1-4, and if it was positive after the reversal the turn was annotated as Ventral, and Dorsal otherwise. These annotations were added to the ethogram (Fig. S3D). For visualization purposes, the turn annotation was extended in Fig. 1 to a minimum duration of 10 volumes (about 3.5 seconds) in order to be clearly visible.

Self-collision is defined by measuring when the nose came close to any other body segment. However, the exact distances can be noisy especially when the body is in a complex shape, so several time series were averaged to stabilize the calculation. Specifically, the pairwise distances between the 4th kymograph segment (the “nose”) and all other segments were calculated. Several quantiles were calculated (0.01, 0.1, and 0.2) and averaged. This time series of close distances was z-scored and smoothed using a rolling mean (window of 5). Finally, it was discretized using a threshold of −2.5, i.e. any time the nose is 2.5 standard deviations closer to any segment than average. These collisions often occurred at the beginning of post backward crawling turns (See example in Fig. S3D).

### Fictive Discrete Behavior Annotations in Immobilized Recordings

For immobilized recordings, behavioral states were annotated as previously reported^1,28,60^. First, the rise, high, fall, and low periods of the AVA trace were annotated (left-right pooled). This was done in the following way:

1. Find the peaks in the derivative in two steps:
2. Find peaks in a strongly smoothed signal
3. Find peaks in the original signal
4. Keep peaks that are in both (or close)
5. Remove peaks that do not have a large enough delta in the original signal
6. Repeat step 1 for negative derivative
7. Assign the positive peak regions as “rise” and the negative peak regions as “fall”
8. Assign intermediate regions based on two passes:
9. If it is after a rise and before a fall and the amplitude is > high_assignment_threshold, it is “high”
10. If it is after a fall and before a rise and the amplitude is < high_assignment_threshold, it is “low”
11. Otherwise it is “ambiguous” and assigned based on the mean amplitude (closer to average of previously assigned high or low)

Then the states are defined as follows:

1. The start of an AVA rise is the start of a Reversal.
2. AVA fall ends the Reversal, and starts a Turn. Dorsal or Ventral identity is defined by comparing the peak amplitude of SMDD to SMDV. Higher SMDD activity is a Dorsal turn, and higher SMDV activity is a Ventral turn. If SMDV is not identified, then RIV is used. All neurons are left/right pooled.
3. AVA low defines the Forward state.

This heuristic was validated by the activities of these neurons individually recorded in freely crawling worms, which served as reliable indicators if the respective behaviors^1^.

### Image Preprocessing

To correct for movement between different z slices of a single volume, each slice is aligned rigidly to the middle slice using the OpenCV package in python.

### Neuron Segmentation

We used a custom-trained Stardist^55,56^ network for 3d segmentation. We found that edge effects generate significant false positives, and thus the network was not applied to the full volume. Rather, a bounding box was calculated in 3d on the entire smoothed image, using a binarization of a very strong Gaussian filter (sigma=8). Segmentation was applied on this portion of the image, significantly increasing speed and accuracy.

### Neuron Tracking

We tracked using a two-step procedure. In parallel, short-time tracklets and time-independent global tracks were calculated. Final tracks were produced by combining these together. Each step is explained in more detail below, and uses a neural network trained on manually corrected tracks of several videos. Two neural networks were used, both modifications of the SuperGlue network^58^. They will be referred to using SG_tracklet_ and SG_global_ for tracklets and time-independent tracking respectively.

Both steps operate on all objects detected by the segmentation algorithm, embedded in a feature space. All individual steps of both tracking algorithms perform set-matching. That is, they operate on two unordered lists of detected neurons, and produce a match with confidence.

### Feature space embedding and neural network training

Each neuron is embedded independently using pretrained feature generators. Specifically, for each z slice from −3 to +3 slices from the centroid of the neuron, we generated a feature vector of length 128 using the OpenCV VGG pretrained neural network. The entire feature vector is a concatenation of these features and the physical position of the neuron.

SG_global_ and SG_tracklet_ were both trained on this feature space, and differ only in the presentation of the training data. SG_global_ was trained on randomly selected pairs of frames throughout the video, whereas SG_tracklet_ was trained only on adjacent time points.

### Time-independent tracking

Using the embeddings above, the time independent or global tracking proceeded as follows:

1. Select 10 random reference volumes (time points).
2. Perform all pairwise matches using SG_global_ within reference volumes and cluster the resulting graph. This generates consistent labels within the reference group.
3. For each reference volume, match all other volumes using SG_global_. This assigns 10 labels with confidence to all other neurons, equivalent to a bipartite graph.
4. Filter edges (matches) in the bipartite graph with unrealistic z changes or low confidence.
5. Perform bipartite matching to generate a final label.

### Tracklets

Using the same embeddings, the short-time tracking proceeds as follows for all time-adjacent pairs:

1. Form a bipartite graph using an ensemble of methods:

a. SG_tracklet_
b. Affine matching
c. fDNC^113^
2. Filter edges (matches) in the bipartite graph with unrealistic z changes or low confidence.
3. Perform bipartite matching to generate a final label.
4. Starting at t=0, extend pairwise matches into tracklets. Tracklets end when there is no match at a time point.

Note that by default and for the datasets in this paper, SG_tracklet_ is the only method used. The others are included for comparative purposes only. The Affine matching is best when there is very little motion, for example during quiescence or to track immobilized animals. We found that the fDNC method was good for certain neuron classes but not the majority, although this may vary by genotypes used in different labs.

### Combining tracklets and global tracks

The global tracks have outliers, and the tracklets can improve them as shown in Fig. S1. Tracklets are assigned in the following way:

1. Build a bipartite graph between true labels and all tracklets. Edge strength is determined by the fraction of each global labels that neurons within a tracklets were assigned by the global tracker.
2. Perform greedy matching to get an initial label-tracklet matching
3. Resolve conflicts. If more than one tracklet is assigned at the same time to the same ID, split them into 2 new tracklets and redo greedy matching
4. Produce the final tracks by concatenating all assigned tracklets

Note that if there is a time point and neuron for which a tracklet is not assigned, then that time point will have a gap in the final tracks regardless of the assignment of the global tracker. Note also that there are many unassigned tracklets, for two possible reasons: 1) the global tracker was too noisy and thus the tracklet could not be matched; or 2) the tracklet itself was too noisy, and could not be matched. When manually correcting tracking, assigning and correcting tracklets is the main task, and is supported in our custom GUI. We found this to be orders of magnitude faster than single-volume track correction.

### Postprocessing tracks

Next, tracking errors in individual neurons were detected using probabilistic PCA. This relies on the intuition that each object should maintain approximately the same distances to all of its neighbors, and that large jumps are due to tracking errors. This can be quantified by calculating all pairwise distances between all objects, and doing a low-dimensional projection to detect outliers. It is important to reduce the impact of far-away neurons in this calculation, and we found a simple weighting by standard deviation of the pairwise distance worked well. This algorithm works well when there are only a few outliers, and performance degrades rapidly for objects with many mistakes.

### Neuron Identification

Neuronal cell identity determination (IDing) was performed using a heuristic that was previously used in refs.^1,28,60,61^; these studies included additional validations using cell specific reporter constructs and / or the NeuroPal method^62^. In general, neurons were identified based on their activity (e.g. correlation to activity of other community members), shape, and relative position. The neuronal identification process starts with identifying the AVA neurons, which are the brightest reversal active neurons located anterior in the head ganglion. Their activity provides a reference for identifying the rest of the neurons. We compared the activity of the same neurons in the same individual in both freely moving and immobilized conditions (n=7) and found that many neurons keep their general activity pattern., e.g., correlation to AVA between these conditions. Therefore knowledge from previous works on immobilized animals is applicable. To validate the gas-sensitive neuronal identities, we exposed the immobilized individuals to 11% to 21% oxygen shifts. As an orthogonal approach, and similar to^60^, we further confirmed cell identities in the head ganglia using NeuroPAL**??**. Specifically, we recorded both NeuroPAL stacks and immobilized whole-brain activity from the same individuals expressing a pan-neuronal GCaMP6f (n=6). We then validated the anatomical position, shape and activitypatterns of the identified neurons. Table S3 lists how each identified neuron is detected, based on the parameters described above. We also provide a confidence level for each ID. The highest confidence level 3 is reserved for neurons that can be identified based on their position and shape alone. Confidence levels 0 to 2 refer to neurons that are identified also using their stereotypical activity pattern. Neurons with confidence level of 0 are uncertain.

### Preprocessing traces

The traces contain some artifacts and noise, qualitatively more than prior immobilized or singleneuron recordings^1,28^.

The traces were initially calculated using the following formula:

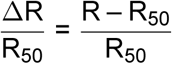

Where R is the ratio of the sum of the background subtracted and bleach corrected red and green channels for each segmented mask, R_50_ is the median of R for a full tracked object. For analysis of sensory neurons with more infrequent responses, R_20_ (the 20th percentile of the trace) is used instead of R_50_. Bleach correction was performed using a direct exponential fit using the lmfit package in Python, and dividing the trace by the model fit. To preserve the relative magnitudes of the green and red channels, the original mean was added back in after dividing by the exponential.

Next, basic preprocessing and filtering was applied to individual traces. Outlier activity time points were initially removed with a threshold of 5 standard deviations from the mean. Traces were then filtered with a rolling mean of window size 5 (≈ 1.5 seconds), which filled-in small gaps of missing data.

Next, further preprocessing steps were applied across the entire dataset. Objects were dropped entirely if they did not meet a tracking threshold, set at 75% of the video. Next, trace values were set to nan if the behavioral tracking of the animal failed or if the frame was corrupted. This was detected by two outlier detection algorithms applied to the number of segmented objects in a volume: LocalOutlierFactor with contamination=0.1 from the sklearn package in Python, as well as a simple threshold decrease in the number of objects detected (set at 40).

Finally, several steps (notably PCA) required dense matrices with no missing values, so any missing values, especially those added in the above steps, were filled in. First, data was imputed using a simple rolling mean with window=3 (1 second). Any remaining gaps were imputed using linear interpolation in the pandas library in Python.

### Categorizing intrinsic activity

#### Comparing PCA weights across individuals

PCA decomposes a data matrix into a set of time series (”modes” or “components”) and a set of neuronal coefficients (”weights”). The weight vectors are sign-indeterminate meaning that when the algorithm is rerun, all weights could have a flipped sign and the corresponding mode will be flipped as well. To compare PCA weights across individuals (Fig. 2C), the signs of the time series were flipped such that it is anticorrelated with velocity.

In all calculations the traces were not z-scored, such that PCA was calculated on the covariance matrix, not the correlation matrix. The PCA algorithm used was the implementation in the Python package of sklearn^114^, and requires a full matrix (no nan values). Thus, any calculation of PCA modes required any gaps to be filled, as described above.

#### Definition of Intrinsic activity

Here we define “intrinsic” activity as activity that occurs in the absence of motor execution hence, without time-varying proprioception, and which does not require acutely fluctuating external sensory stimuli. These conditions are met upon our immobilization procedure, where animals do not move their bodies, and any other environmental conditions such as light exposure, temperature and O_2_ concentrations, are kept constant. We conclude that features in neuronal activity that are shared between immobilized and freely moving conditions do not require such inputs and most likely are driven by intrinsic factors, such as spontaneous circuit dynamics, central pattern generators and connectivity. Here, we focus on the covariance structure as a feature of neuronal activity, which results from frequent transitions in neuronal activities and reproducible similarity thereof across many neurons. This can be quantified using the weights of the first PCA component. There are four properties that determine the categories shown in Fig. 2C-D: 1) The distributions are significantly different from each other; 2-3) either distribution is significantly different from 0; 4) the sign of the median of the weights is the same between conditions. The categories are then defined in the following way:

1. If the freely moving condition is not significantly different from 0, then it is “No manifold.”
2. Otherwise, if the difference between the two conditions is significant, then it is “Intrinsic” if the sign of the medians is the same and “Encoding switches” (no cases found) if the sign is different.
3. If the immobilized is not significantly different from 0, but the difference is significant, then it is “Freely moving only”.
4. Finally, regardless of the 0-comparison significance of the immobilized condition, if the difference is not significant, then it is “Intrinsic”; this forms the majority of “Intrisic” category neurons.

Thus, “intrinsic” categories neurons are mostly neurons with no significant difference between their weight distributions. However, some, like AVA, have a significant difference between the distributions but do not switch signs. This change in weights for AVA is not due to a significant change in the qualitative time series of AVA, but seems to be due to the relative balance of strong activity neurons which form PC1 in the two conditions. Specifically, in the freely moving condition AVA is very high signal-to-noise and dominates PC1 (which can be seen in the weights in Fig. 2C), whereas in the immobilized condition there are a larger number of high signal-to-noise neurons, reducing the dominance of AVA.

There are several limitations of this method, especially that it only uses PC1. Thus, activity in higher PC modes may be very different. However, directly extending this method to higher modes is difficult, because the interpretation of higher modes is not consistent across individuals (examples in Fig. S6). Future work would likely need to use more complex manifold alignment methodology.

We cannot exclude environmental conditions that arouse the animals and thereby promote the dynamics we observe, like laser-excitation light^68^, the recent experiences of food removal^49,115^, and/or the prior transfer to the assay arena^116^. However, such stimuli do not exhibit any temporal patterns that could align to the transition patterns we observe in our data. Therefore, exafferent inputs are highly unlikely to be major immediate causes of the observed rapid switches in neuronal activity. In addition, it is important to note that it is easier to identify neurons with such correlations, therefore Fig. 2D describes an upper estimate on intrinsic activity.

#### Significance calculations

Unless otherwise stated, t-tests are calculated using the scipy library^108^ in Python, using the ttest_ind or ttest_rel (for paired samples) implementation. For clipped input statistics like PC1 weights (Fig. 2C), the ttest is calculated via bootstrapping, not the t-student distribution (i.e. setting the argument “permutations=100000”).

Correction for multiple comparison is done using the ‘fdr_bh’ implementation of false discovery rate in the statsmodels library^117^ in Python.

#### Quantifying accuracy of the computational pipeline

The accuracy of the pipeline was calculated using manually annotated ground truth test dataset, not used for training the neural networks. Adding multiple reference volumes improves the median accuracy, but only a small percentage for many neurons. Across neurons, the majority of tracked objects are further improved by adding tracklets, which is the full pipeline.

Accuracy was calculated on a held-out dataset not used for training the tracking or segmentation networks. Note that not all objects could be manually corrected, and thus all metrics are calculated on a subset of ground truth annotated objects.

#### Motion Artifacts

There are considerably more motion artifacts in freely moving animals when compared to immobilized ones, namely nonrigid deformations and stretching. In addition, there is image tearing across a single volume when the animal is moving particularly quickly.

Although a full quantification of all possible artifacts is beyond the scope of this work, we tried to carefully validate the low amplitude residual signals that we analyzed in Fig. 3. We did this in three ways:

1. Identifying the same neurons when possible in GFP recordings (which display only motion artifacts), and confirming that the behavioral encoding (specifically, oscillations), was significantly different. This can be seen by the (Fig. 3I) separation between VB02 and other motor neurons (RMED/V) in GFP (gray) and in GCaMP (red).
2. Identifying neurons with similar positions but opposite oscillations. This is important in particular because oscillatory activity could be due to artifacts correlated to body bending, which should be shared by neurons in the same region and side of the body. Specifically, VB02 and DB01 display ventral and dorsal oscillations respectively (Fig. 3E,F,K), even though they are very close within the Ventral Nerve Cord and both on the ventral side. This is consistent with their opposite innervation patterns and proposed functions: VB02 innervates ventral body wall muscles and is thought to drive ventral body bending, whereas DB01 innervates dorsal body wall muscles is thought to drive dorsal bends^80,118,119^.
3. Consistency of behavioral encoding across individuals. Specifically, the coefficients in the Bayesian model for the behavioral encoding were shared across individuals, reducing the possibility of overfitting noise in any one dataset. We found this was particularly important for certain motor neurons like RMED/V, which display correlations to various noisy kymograph segments at the individual dataset level in GFP, but are not consistent across recordings.

#### GCaMP testing

We found that there were dramatic differences in signal between different GCaMP lines, such that our previous line of GCaMP6f used in prior immobilized whole-brain and freely moving single neuron recordings^1,28,60^ was insufficiently bright under these conditions and showed very low signal to noise. In that line, fewer than 10 neurons were clearly active on average, and many recordings showed no obvious activity (data not shown). For this reason we tested several GCaMP lines (6f, 7f, and 7b), and empirically determined that 7b showed the most clear activity in our conditions.

### Computational analysis

#### Canonical Correlation Analysis

We used Canonical Correlation Analysis (CCA), a dimensionality reduction technique that builds a linearly optimal shared subspace between two input datasets, in this case the top 5 PC modes of neural dynamics and selected behavioral time series. PCA modes were used for two reasons. First, they are the main object of study in this work. Second, interpreting the weights of linear regression methods is difficult when input variables are correlated^120^. In this case, many neuronal time series are highly correlated, and could receive opposite or highly variable weights in a naive linear regression. Preprocessing the data to remove correlations, such as via PCA, is one way to stabilize the final neuronal weights, and these are the CCA weights that are shown in Figs. 1 and S5.

#### Triggered Averages

For all triggered averages, a set number of preceding (40 volumes, 12 seconds) and subsequent (100 volumes, 30 seconds) time points were extracted and organized into trials. For event triggered averages, time points were removed from the preceding period if another instance of the triggered event occurred, and similarly any data points after the end of the triggered event were removed. Then, trials were averaged within one dataset. The mean and standard deviation shown in Figures 3, 4, S8, and S7 are taken with respect to datasets.

#### Power Spectra

Power is calculated using Welch’s method^121^ in the Python package Scipy^108^, which denoises the calculation by taking the average of the power spectrum across multiple overlapping windows.

#### Bayesian modeling for hierarchy quantifications

We performed multilevel Bayesian modeling using the PyMC package^122^ in Python. Such Bayesian analysis in neuroscience has several benefits over traditional least-squares model fitting [123], and we describe the most relevant ones below. First, it generates a distribution of model parameters given the data, instead of a single best value. This helps prevent overinterpretation of parameters whose average value may seem interesting, but whose variation is very large or overlaps with 0. The second main benefit is the ability to fit multi-level models, in which parameters nest within one another. This is naturally applicable to our experimental data, for example if we want to fit a behavior to neuronal activity from multiple individuals (multiple repeats). Assuming our model structure is good, we believe that there is an underlying cross-individual parameter describing the relationship between the behavior and the neuronal activity. However, there can also be inter-individual differences, which we want to model separately from the underlying cross-individual parameter. The output a multi-level organization is a separate distribution of parameters per individual (lower level), as well as a parent distribution of the underlying cross-individual parameter (higher level). Third, Bayesian analysis explicitly models the residual noise distribution, i.e. the variation in the data that is not explained by the core model terms. This is particularly useful when there are outliers in the data, as the standard least-squares fitting procedures can be dramatically influenced by them.

Another practical point in Bayesian analysis is the specification of prior distributions, which can regularize models away from impossible values and focus on realistic regions of parameter space. In this work, we fit three models to the combined neural traces from multiple individuals, using a multi-level approach. The multi-level aspect is highlighted by hyper priors, which are prior distributions that describe a variable which is itself transformed into another distribution. For example (as described below) we can have a hyperprior for a dataset-independent offset in the data, which is then a variable in a prior describing a dataset-dependent offset. This is especially useful for focusing on the value of parameters that are dataset-independent, while still modeling the differences between datasets.

See Table S1 for number of datasets per neuron.

#### Null model

First, as a control for the subsequent models, we fit a null model which does not use behavior information to predict neuronal traces. The core model is:

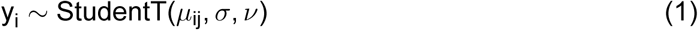

where y_i_ is the neuronal trace to be explained, *µ*_ij_ is the mean predicted value, in this case a constant offset (different per neuron and dataset). Further, i indexes the neuron, j indexes the dataset, and StudentT describes the residual term, modeled as a StudentT distribution. This is a distribution that is more robust to outliers than the classic normal distribution. It has two parameters, *σ* and *ν*, which each have prior distributions. Along with the prior distribution for the constant parameter, there are 6 additional equations to specify the full null model:

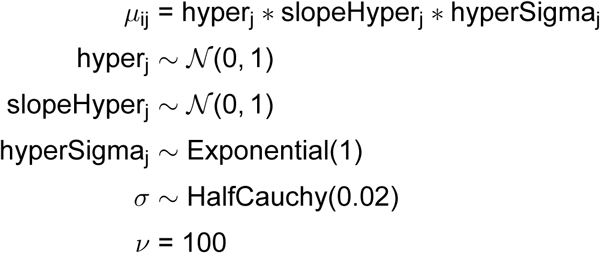

where *µ*_ij_ is a described by “hyper-priors” that allow values to be different per dataset. These hyperpriors have “hyper” in the name (hyper_j_, slopeHyper_j_, and hyperSigma_j_), where N(0, 1) is the normal distribution centered at 0 with unit variance. Note that *µ*_ij_ is expanded into a z-scored version, which is a standard technical detail that improves numerical convergence. The HalfCauchy and Exponential distributions are standard prior for parameters that must be positive, such as variances. All default values were determined using prior predictive checks, where time series were simulated using the model parameters and were checked to make sure they are reasonable. In addition, = refers to a simple deterministic equation, and ∼ refers to a random variable which is drawn from a distribution, and for which a posterior distribution is fit using data.

#### Behavior model

Each model has the same core form as Equation 1, with the same priors on *σ* and *ν*; the difference is in the prediction of *µ*_ij_, i.e. the mean of the model. Instead of a constant *µ*_ij_, the second model uses behavior as a regressor to predict the time series. To make the notation cleaner, the subscript i will be dropped, as all neurons are modeled independently, but j will be kept to show which parameters are allowed to vary across datasets:

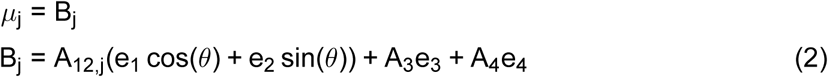

where B_j_ is the combined behavioral predictor term, the e_j_ components refer to eigenworms, and the coefficients A_12_, A_3_, and A_4_ are the coefficients for the eigenworms. The first two eigenworms are converted to polar coordinates, thus they have a combined amplitude (A_12_) and an angle (*θ*). The phase (*θ*) is plotted in Fig. 3K when fit using the third model, described below. Each dataset is allowed to have a different eigenworm 1 and 2 amplitude, however, the encoding of other eigenworms was not varied across datasets. This is done for both interpretability as a well-defined phase shift across all individuals, as well as allowing the amplitude alone (A_12_) to vary across datasets. Thus there are hyperpriors only for A_12,j_:

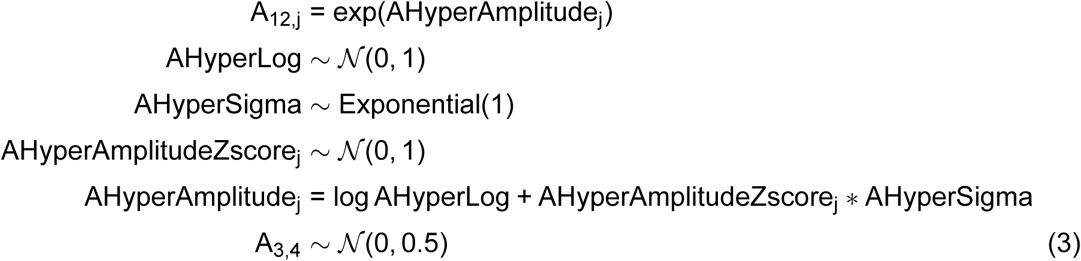

where the parameter for A_12,j_ has been expanded in a zscored version and calculated in log space, both technical details that improve numerical convergence. In more detail, AHyperLog does not vary across datasets and captures the amplitude independent of dataset, and AHyperSigma captures a scale parameter independent of dataset. AHyperAmplitudeZscore_j_ is dependent on the dataset, and captures the dataset-specific variation in the amplitude of the final parameter, A_12,j_. The dataset-independent versions of the coefficients are shown in Figs. 3J,L and S7N,O, when fit using the hierarchical model described below.

#### Hierarchical model

The third model (the hierarchical model) contains a single additional term, multiplying the behavior term per time-point by a sigmoid. This allows the relationship between behavior and neuronal trace to vary across time:

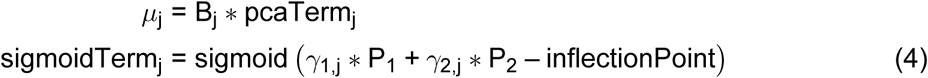

where sigmoidTerm_j_ is a sigmoid function applied to a weighted sum of the first two PCA modes, P_1_ and P_2_. Note that if the input to the sigmoid function is always 0, then the function returns a constant, corresponding to weak or no hierarchy. The coefficients *γ*_kj_ (where k refers to pca mode index) of these modes varies between datasets and is defined by a hyperprior, almost exactly like the coefficients of the eigenworms 1 and 2 in Equation 3:

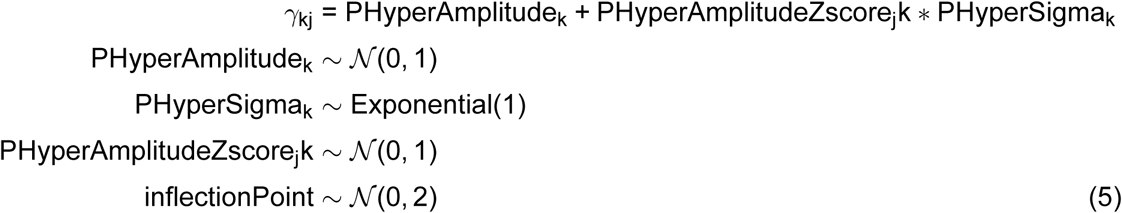

Each of these hyper parameters has a similar meaning as those in Equation 3, but it is not calculated in log space. This is because the amplitude itself can be positive or negative and the parameter for the first mode, PHyperAmplitude_1_, is shown in Fig. 3J and for the second mode, PHyperAmplitude_2_ in S7N. The full distribution of A_3_ is shown in 3L and for A_4_ in S7O. The radial term shown in Fig. 3K is a combination of the dataset-independent eigenworm amplitude in Equation 3 and a representative value for the output of the sigmoid gating term, sigmoidTerm. Specifically, shown is exp AHyperLog ∗(sigmoidTerm, 0.8), using the 80% as the representative value of the non-gated output of the sigmoidTerm.

#### Model comparison

Comparisons between models are derived by calculating the expected log pointwise predictive density (ELPD) using the arviz package^124^. In Fig. 3I, the improvement between the behavior model and the null model (x axis), and the improvement between the hierarchical behavior model and the null model (y axis) are plotted. If the more complex model yielded worse results, a value of 0 was plotted to reduce emphasis on such overfit examples. Further improvement of the hierarchical model over the behavior model is plotted in Fig. S7M (y axis).

### Data and Code Availability

All original code for both video processing and subsequent trace analysis will be available on GitHub upon publication. Data will be available upon publication.

## Acknowledgements

The computational results of this work have been achieved using the Life Science Compute Cluster (LiSC) of the University of Vienna. The authors would like to thank Ev Yemini for the NeuroPal worms and in alphabetical order Netta Cohen, Max Jösch, Harris Kaplan, Meital Oren-Suiza, and Julia Riedl for advice, and Maria Boehm, Hannah Eckert, Sarah Foerster, Eva Gratzl, Leon Lichtenstern, and Anahita Sedighi for manual annotation of tracks. The research leading to these results has received funding from the European Research Council (ERC) under the European Union’s Horizon 2020 research and innovation programme (*elegans*BrainBodyEnvi, #101054527), from the Simons Foundation (#543069 M.Z., NC-GB-CULM00003196-01 M.Z., SFI-AN-SURFiN-00008619 H.B., AN-SURFIN-00003578 H.B.), the International Research Scholar Program by the Wellcome Trust and Howard Hughes Medical Institute (#208565/A/17/Z), the University of Vienna, and the Research Institute of Molecular Pathology (IMP). C.F. and I.L. were supported by VIP2 postdoctoral fellowships co-funded by the European Union’s Horizon 2020 research and innovation programme under the Marie SkłodowskaCurie grant agreement No. 847548. I.L. thanks the support from the EMBO funding agency (ALTF 1037-2019) and the Human Frontier Science Program fellowship (LT000335/2020-L).

## Author Contributions

M.Z., U.R., C.F., and I.L. conceived and designed the experiments. C.F. developed computational methods and analyzed data. I.L. developed the neuronal cell class identification heuristics, performed experiments, and analyzed data. U.R. designed the behavioral arena and the behavioral analysis pipeline, performed experiments, and analyzed data. L.H. designed and built the behavioral arena and microscopy setup. H.B. performed experiments and analyzed data. M.Z. led the studies. M.Z., C.F., and I.L. wrote the manuscript.

## Supplemental Figures

**Figure S1.**
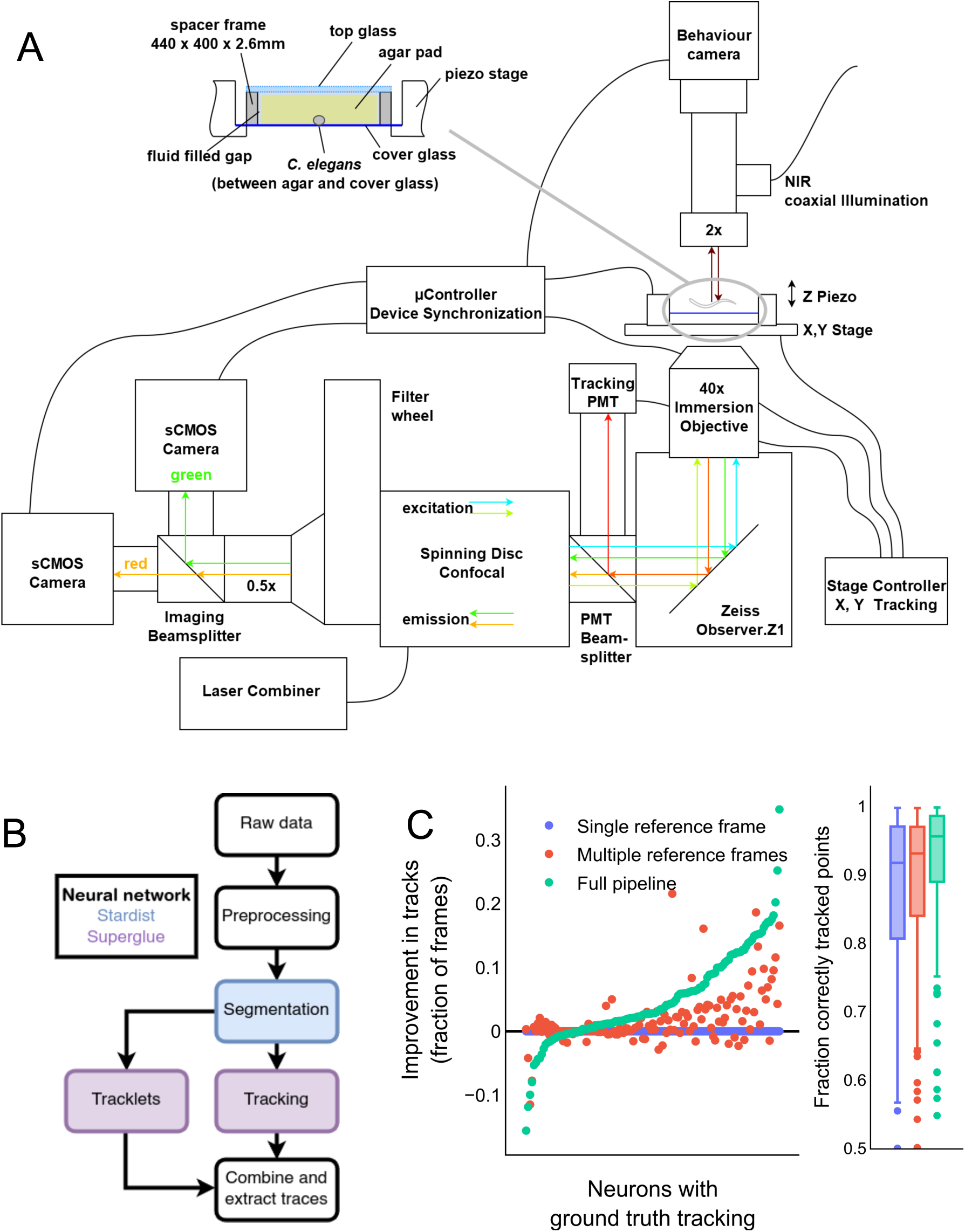
Overview of the hardware and software pipelines. **(A)** Overview of confocal fluorescence and behavioral microscope setup (not to scale). **(B)** Overview of modular computational pipeline, with segmentation on the red channel and tracking performed independently. Tracking is performed in two stages: time-independent tracking and short-time tracklets. See Methods for details on how they are combined. **(C)** Ablation study of individual algorithm elements. Metrics are calculated on a manually tracked test dataset not used for training the neural networks (131 neurons). Boxplots show median, interquartile range, and 1.5 times the interquartile range.

**Figure S2.**
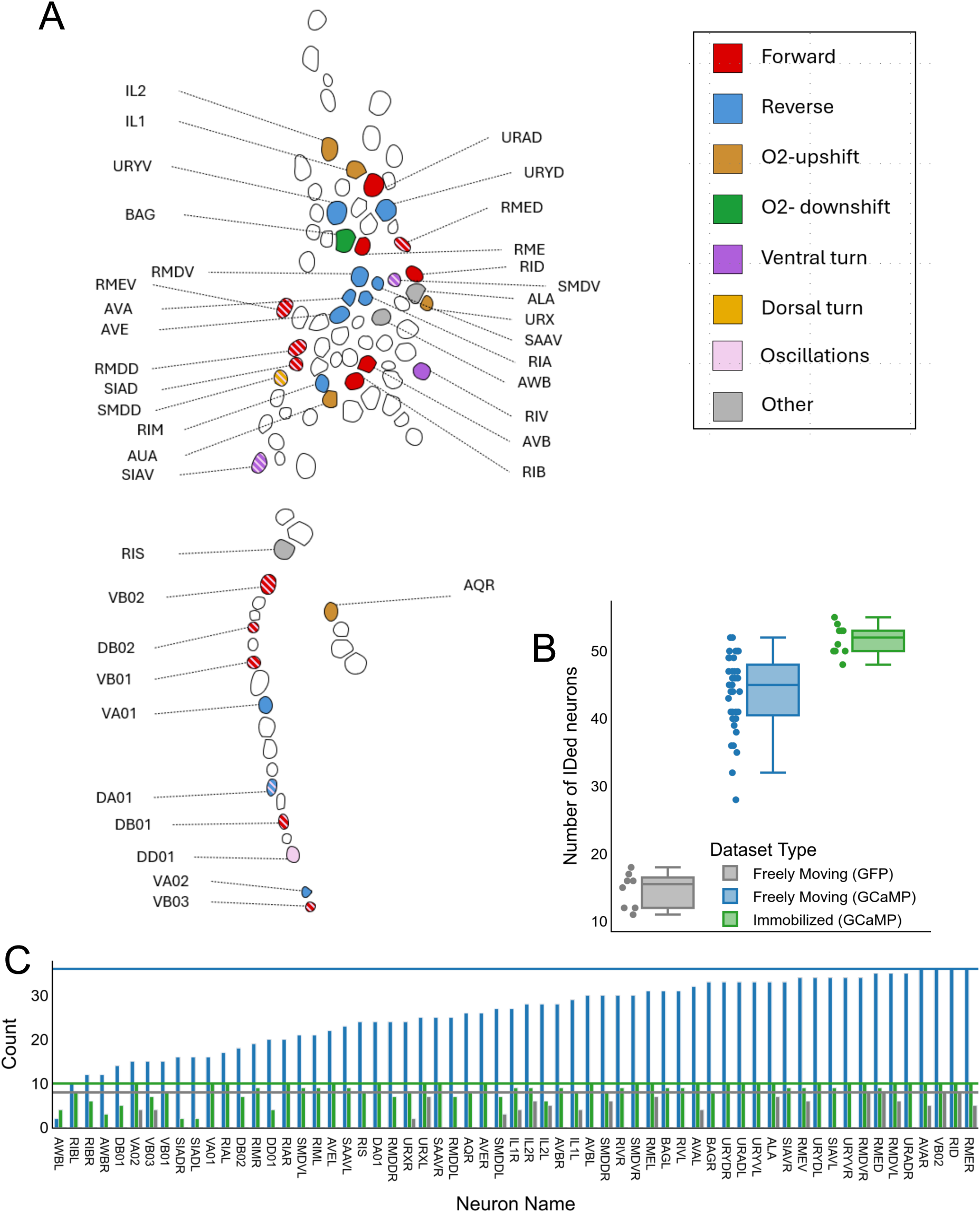
Identification of neurons via position and activity patterns. **(A)** Scheme of neuron nuclei in worm head and ventral cord region. The scheme is based on a maximum intensity projection of one volume from a whole-brain freely moving recording. The typical position and activity patterns of the neurons identified in this work are marked. Neurons are colored by their typical activity pattern: active during forward, reverse or turning movement. Neurons that show oscillatory activity are also colored. Neurons that exhibit two activity patterns, for example forward activity with oscillations, are indicated in stripes of both colors. In addition, O_2_/CO_2_ sensing neurons, showing a typical forward-reverse activity, are colored accordingly. **(B)** Number of neurons identified per dataset. Boxplots show median, interquartile range, and 1.5 times the interquartile range. **(C)** Number of neuron_3_s identified per neuron type and dataset type; same color code as in (B). Horizontal lines indicate the total number of each dataset type.

**Figure S3.**
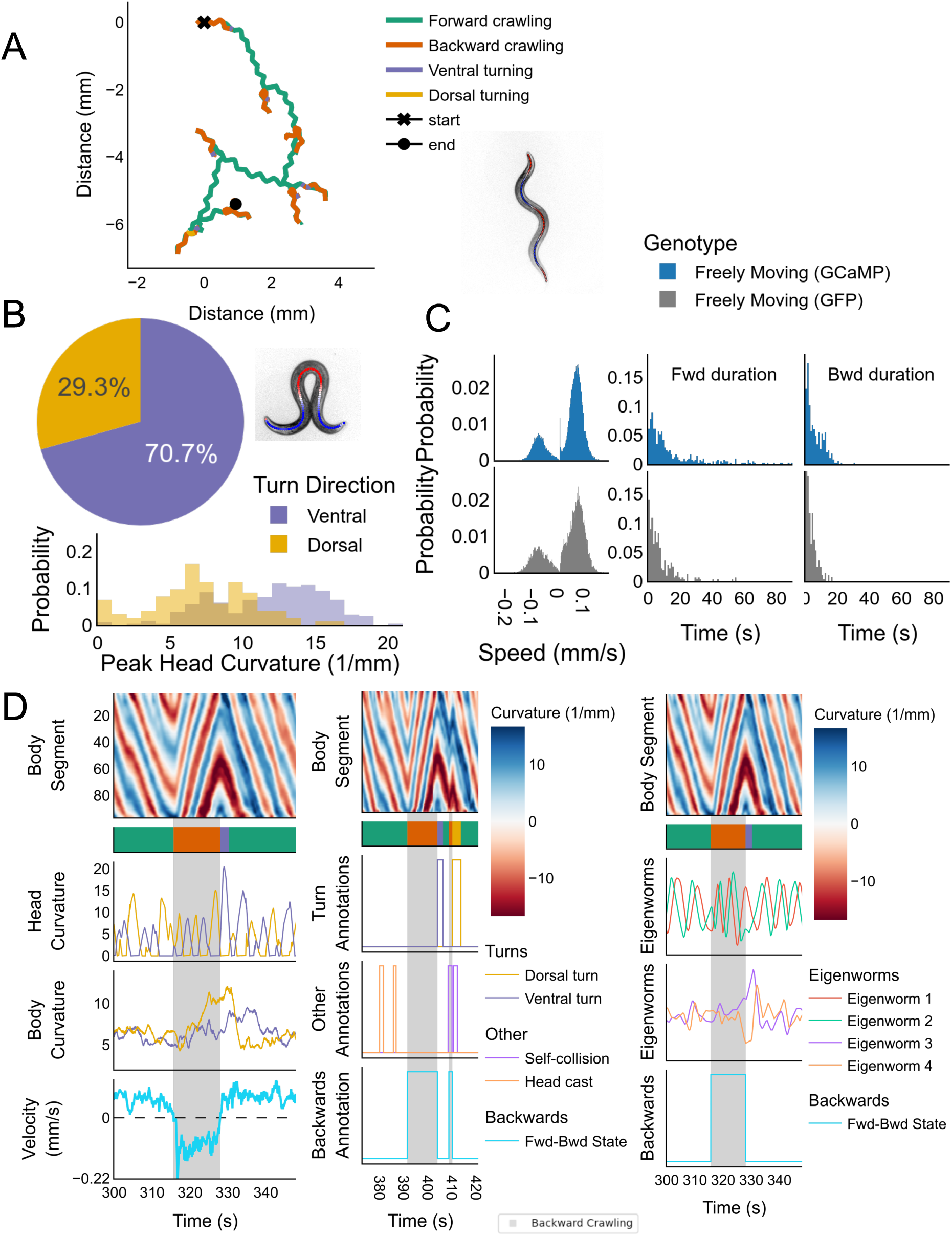
Behavior quantifications. **(A)** Example trajectory of a worm (centroid) exploring an isotropic environment. Further quantification and discrete behavioral labeling is based on the curvature of the centerline shown on worm image in right panel extracted from behavioral videos (See methods). **(B)** Statistics of turn amplitudes across recordings (n=36; bin size=1/mm). An example centerline in an omega turn shape is shown. **(C)** Statistics of speed (bin size=2m/s), forward crawling state duration (bin=1s), and backward crawling state duration (bin=1s) across recordings (n=36 GCaMP, n=8 GFP). **(D)** Three example kymographs (same recording as (A)) of a portion of a recording with their corresponding ethogram (color coded like in (A)). Below the corresponding time series of head and body curvature, worm speed, eigenworm or binary turn, collision, and additional discrete behavioral ann_4_otations are plotted. See methods for definition of each time series or state annotation.

**Figure S4.**
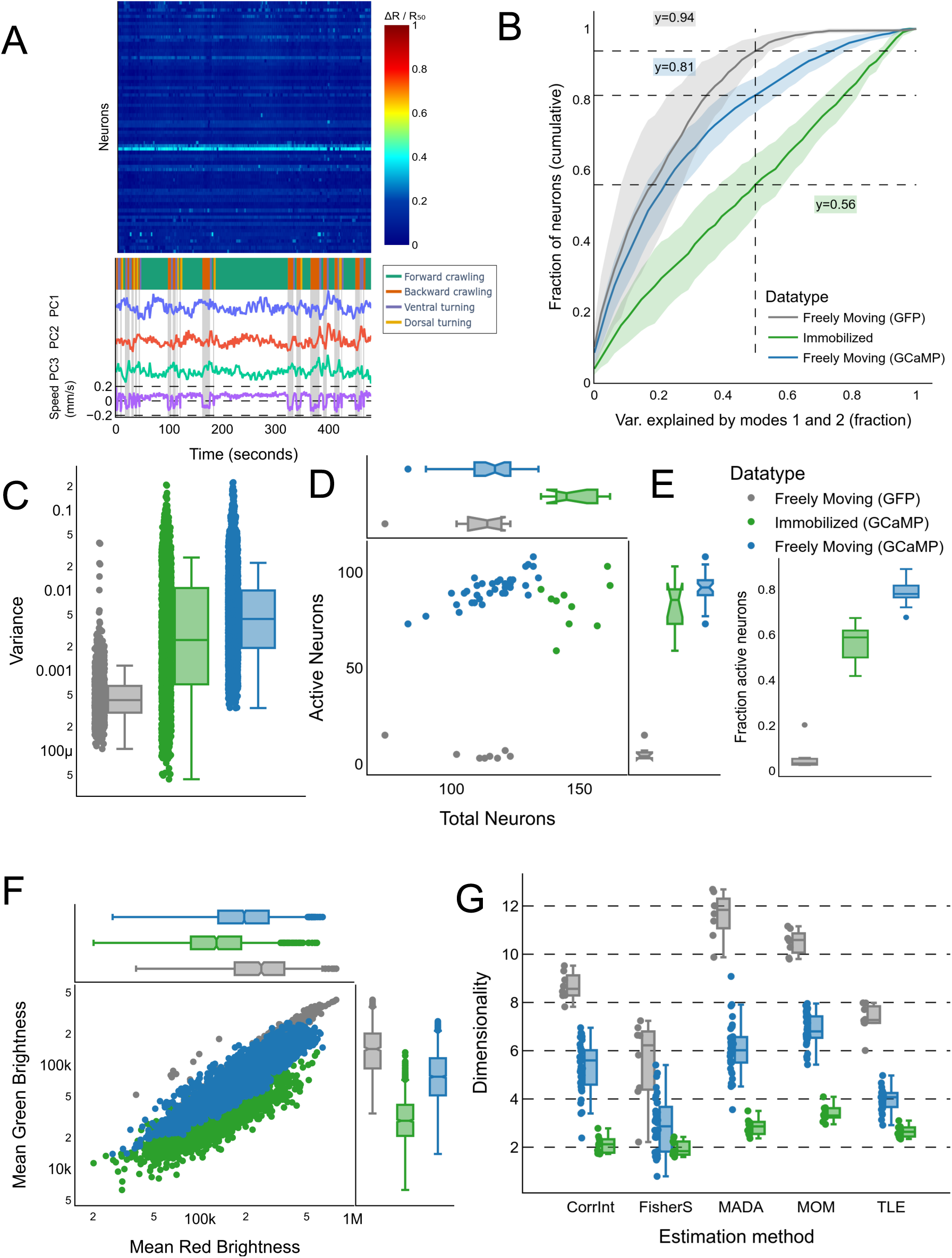
Comparison between GCaMP and control GFP recordings. **(A)** Example GFP recording, which shows no obvious activity. Shown below are the top 3 PCA modes, which show no clear structure. **(B)** The variance explained by the first two PCA modes in immobilized (n=10) and freely moving (n=36) GCaMP and GFP (n=8) recordings. Shown is the mean across datasets plus or minus the standard deviation (shaded region). **(C)** The variance of the mean of ΔR/R50 traces for neurons in all dataset types. **(D)** The number of tracked objects in all dataset types. Objects with less than 75% of time points tracked were discarded. The number of active neurons is defined by neurons with variance above the 95% percentile of all GFP neurons (by definition, 5% of GFP neurons are active). **(E)** The fraction of active neurons in all dataset types, i.e. the ratio of the x and y axes in panel (D). **(F)** The raw intensity values in the green and red channels, averaged across time for all dataset types. Each datapoint represents a tracked neuron, pooled across all recordings. **(G)** Dimensionality of raw data as measured by several different methods. Boxplots in all panels show median, interquartile range, and 1.5 times the interquartile range.

**Figure S5.**
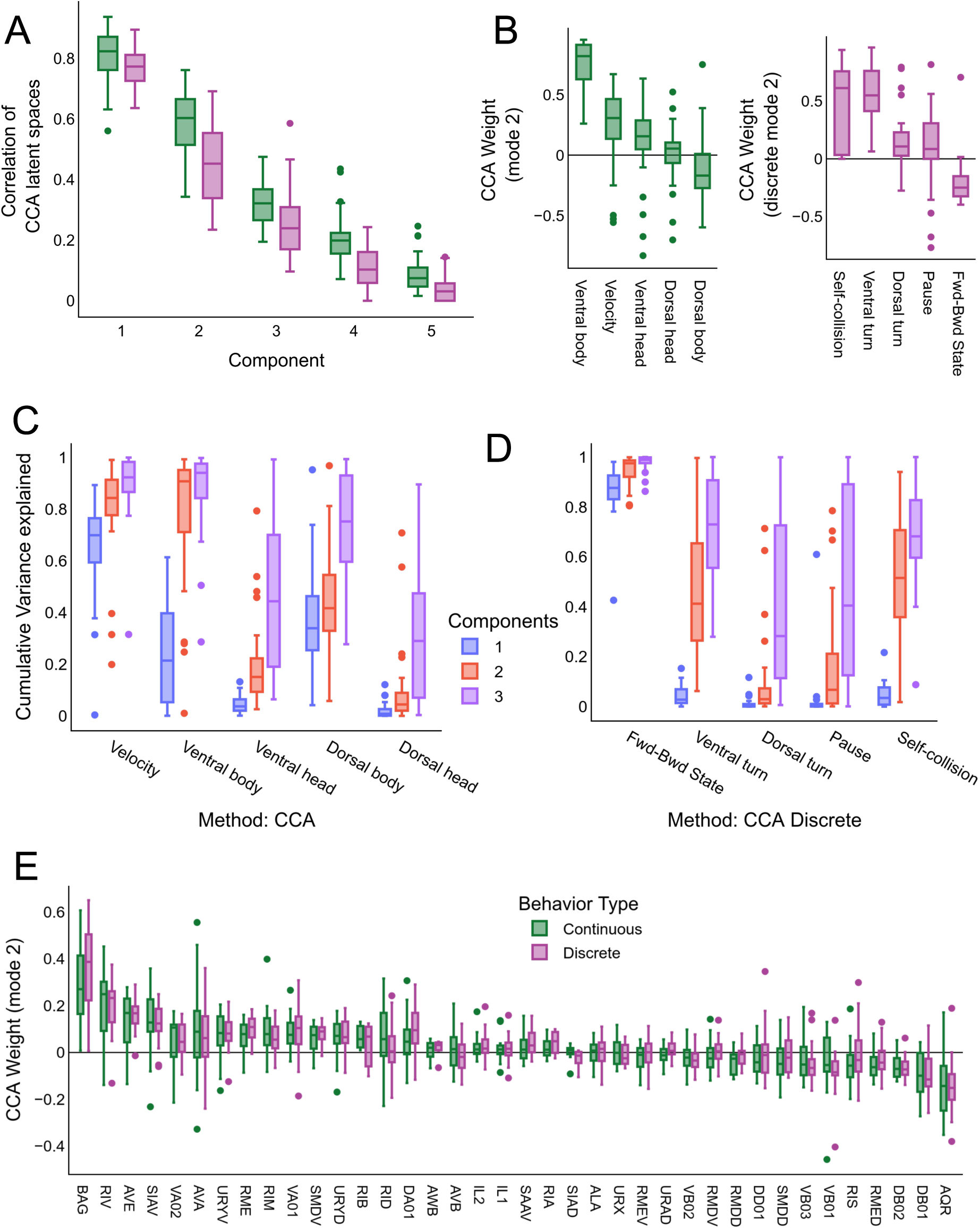
Higher modes of CCA. **(A)** Correlation of latent space generated from neuronal and behavioral data per mode. Ordered with decreasing correlation between neuronal and behavioral data. Same as Fig. 1d but showing additional modes. **(B)** Behavior CCA weights for both continuous and discrete CCA methods. **(C-D)** Cumulative neuronal variance explained by CCA **(C)** or CCA discrete **(D)**. **(E)** Neuronal CCA weights for component 2 for each behavior type, ordered by magnitude in the Continuous behavior model. Boxplots for all panels show median, interquartile range, and 1.5 times the interquartile range.

**Figure S6.**
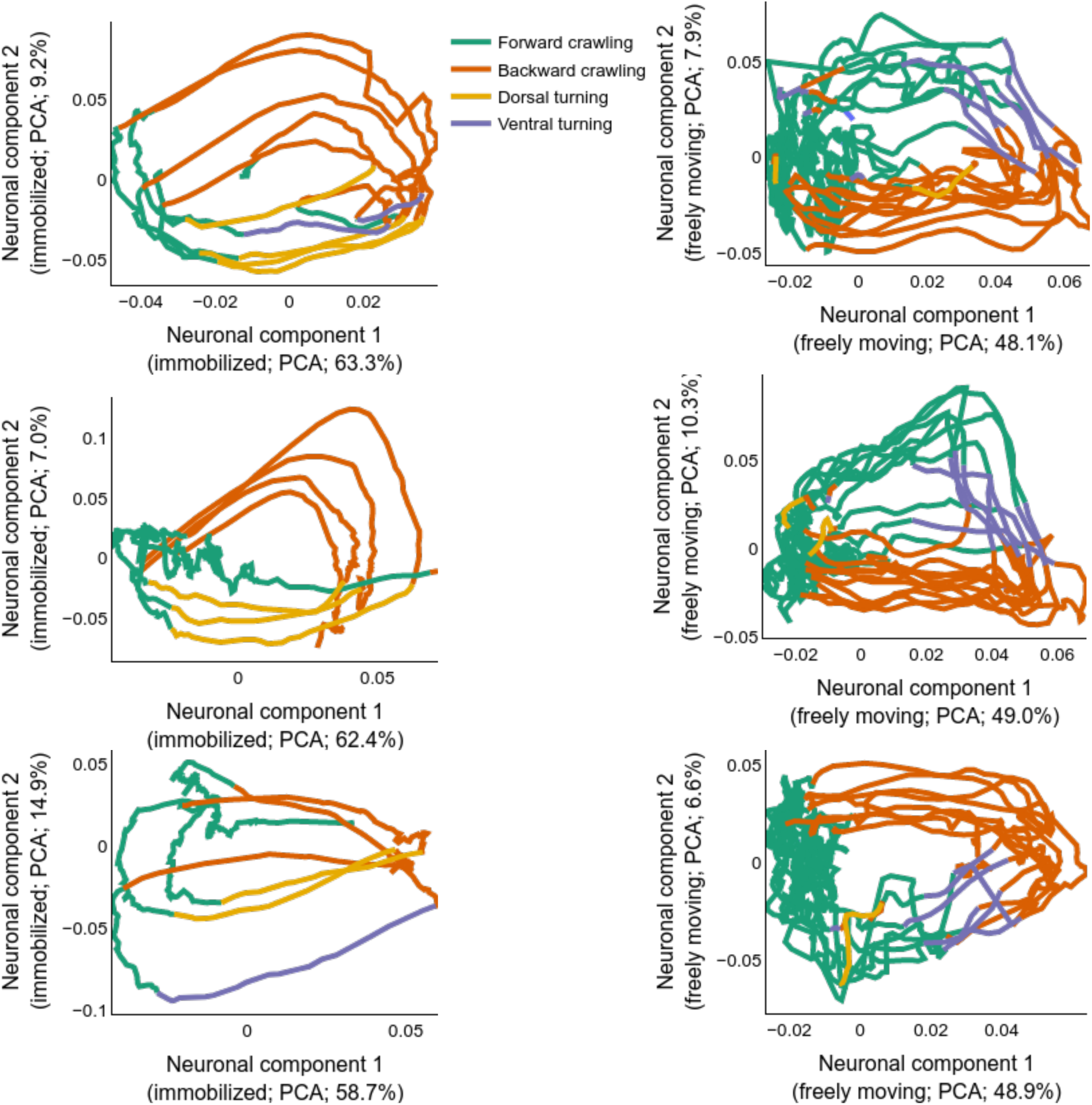
Example PCA manifolds. Example of phase spaces for neuronal activity of immobilized (left) and a freely moving (right) animals. The traces are color coded based on the motor-command state as defined for freely moving or immobilized recordings (see Methods).

**Figure S7.**
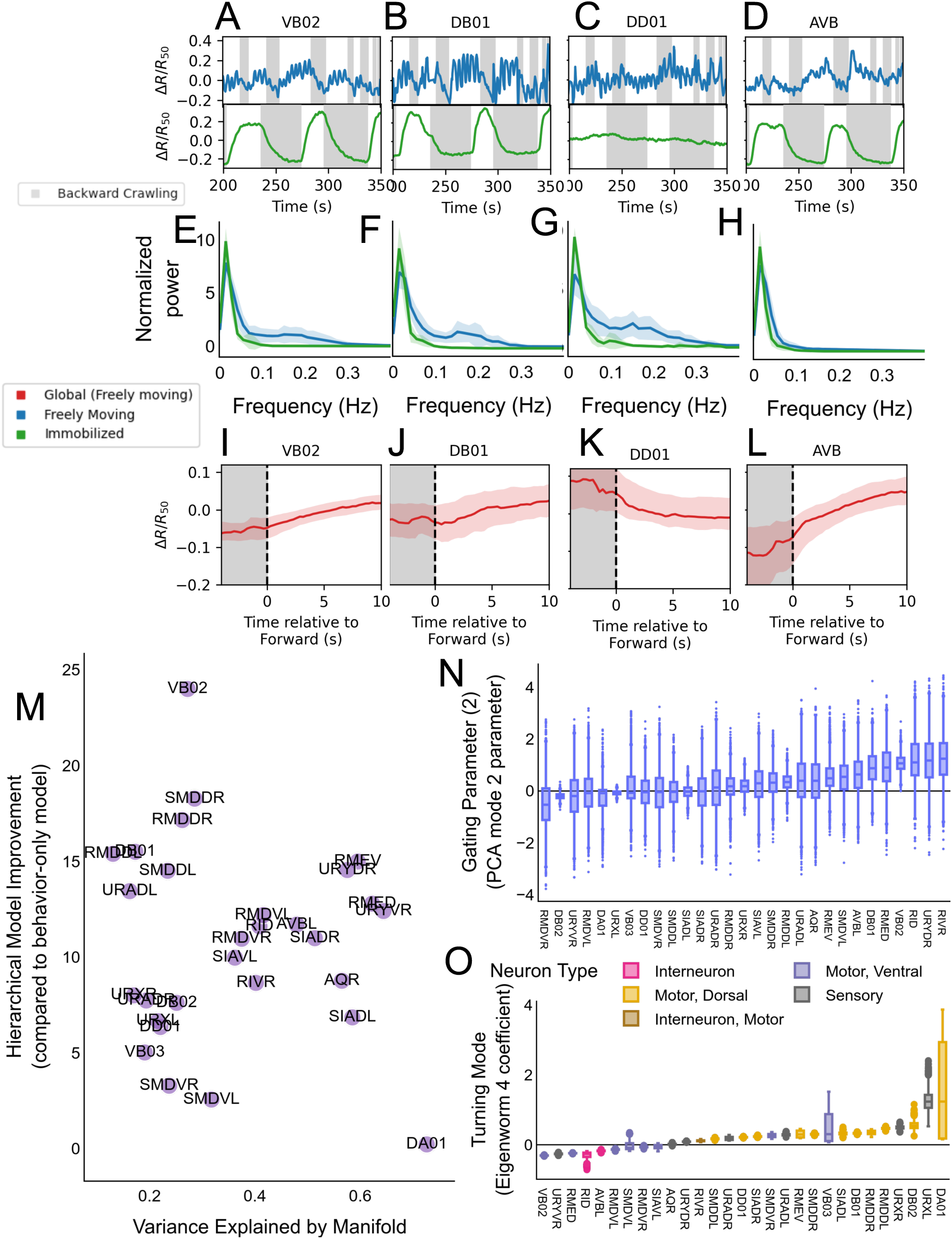
Additional information about individual neurons. **(A-D)** Example time series for the motor neurons VB02 **(A)**, DB01 **(B)**, DD01 **(C)**, and AVB (left/right averaged). Freely moving (blue) and immobilized (green) generally show similar slow components, but there are oscillations in the freely moving condition. **(E-H)** Fourier frequency analysis of neurons across individuals and conditions for the motor neurons, VB02 (**E**; n=36/10), DB01 (**F**; n=18/5), DD01 (**G**; n=19/4), and AVB (**H**; n=35/10). There is a clearly visible frequency peak unique to the freely moving condition just under 0.2 Hz in each neuron except AVB, corresponding to the body undulation frequency. N numbers correspond to datasets in freely moving / immobilized datasets. Figure S7. **(M)** Improvement in model performance for hierarchical model over behavior-only models (y-axis), against variance explained in the raw trace by the manifold (x-axis). Shown are neurons that pass the GFP threshold in Fig. 3i. **(N)** Posterior distribution of the model parameter corresponding PCA component 2 (See Methods). This is also a gating parameter, but does not show very clear structure. PCA component 2 does not have a clear interpretation across animals. **(O)** Posterior distribution of eigenworm 4, which has been shown to correspond to turning behaviors (similar to eigenworm 3). Boxplots show median, interquartile range, and 1.5 times the interquartile range. N numbers for each neuron class are provided in Fig. S2C and Table S1.

**Figure S8.**
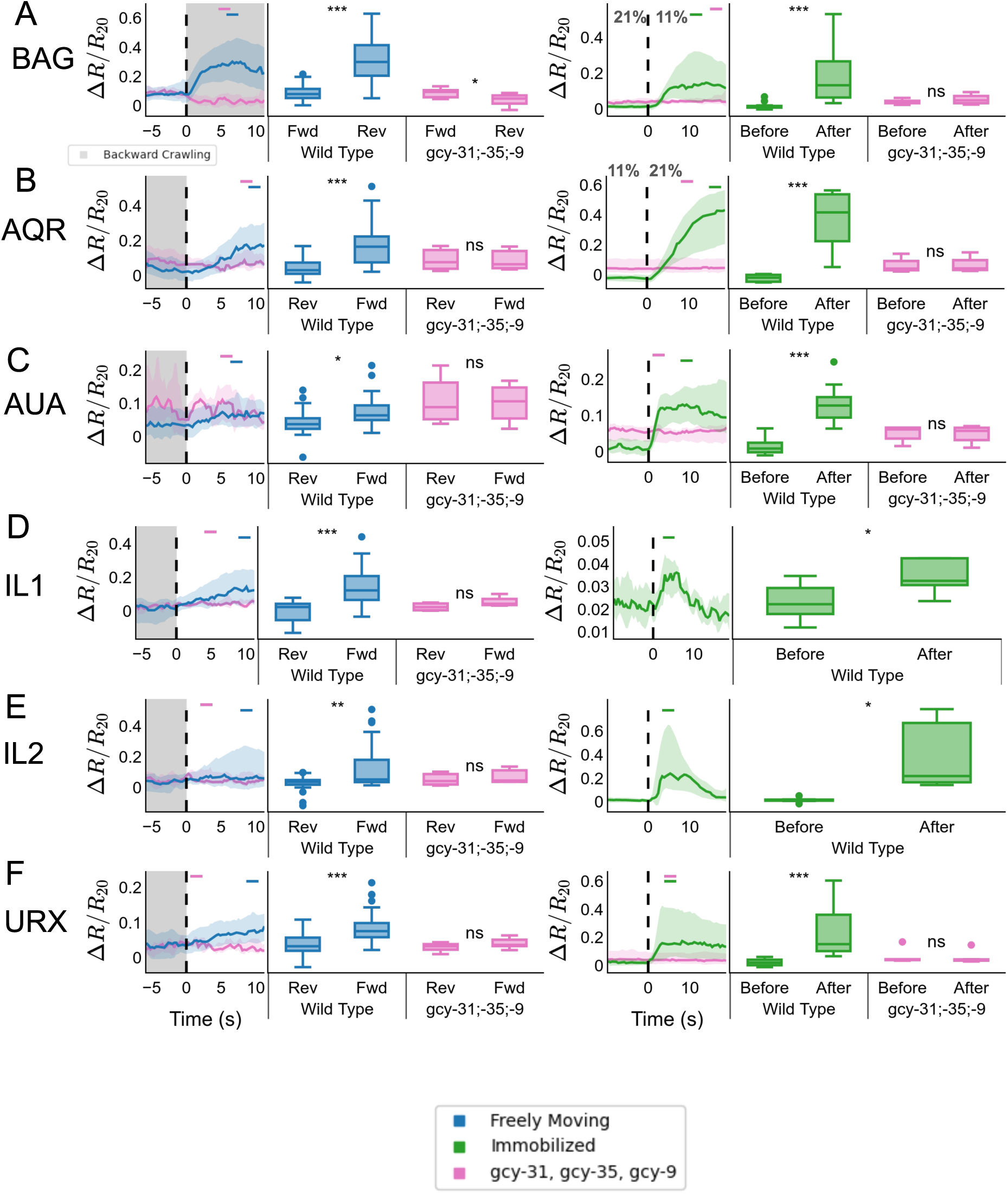
Activity of O_2_ and CO_2_ responding neurons. **(A-F)** First column: triggered averages for wild type and O_2_/CO_2_ sensory mutant (gcy-31;-35;-9) in freely moving datasets. Second column: comparison of preand post-reversal activity in each genotype in freely moving datasets. Third column: triggered averages to O_2_ upshift (BAG is instead shown with respect to downshift) in immobilized animals. For IL1 and IL2 the immobilized recordings were done using histamine-gated chloride channel inhibition of all muscles instead of tetramisole, to allow responses in these neurons. Forth column: comparison of pre- and post-O_2_ upshift (or downshift for BAG) activity in each genotype in immobilized datasets. All traces are L/R averaged (except AQR, which is a single neuron). Shown is the mean across datasets plus or minus the standard deviation (shaded region). See Table S1 for number of identifications and events for all neurons and genotypes. Boxplots show median, interquartile range, and 1.5 times the interquartile range. Significance is shown at the following levels: *, 0.05 < p <= 0.01; **, 0.01 < p <= 0.001, ***, p < 0.001. For all triggered averages, the small bars refer to the time window used for ttest (see Methods).

## Supplemental tables and video

**Supplemental Video 1** Top left: worm undergoing locomotion behavior, viewed with an infrared camera. Frame rate is 3.5 Hz. Top right: 3D PCA projection of the neuronal activity, color coded by action state. Black dot shows current time point. Bottom half: Ethogram of action state (same colors as in the PCA plot), with short states (turns and pause) extended to a minimum of 3 seconds for visibility. Below is a heatmap of neuronal activity across 121 neurons, sorted by PCA mode 1 weight. White line indicates current time point.

**Table S1. IDs of neurons per condition** Excel file containing number of times each neuron was identified in each condition. This includes event numbers for triggered averages

**Table S2. Neurons in intrinsic and other categories** Excel file containing categorizations of all neurons in different conditions.

**Table S3.**
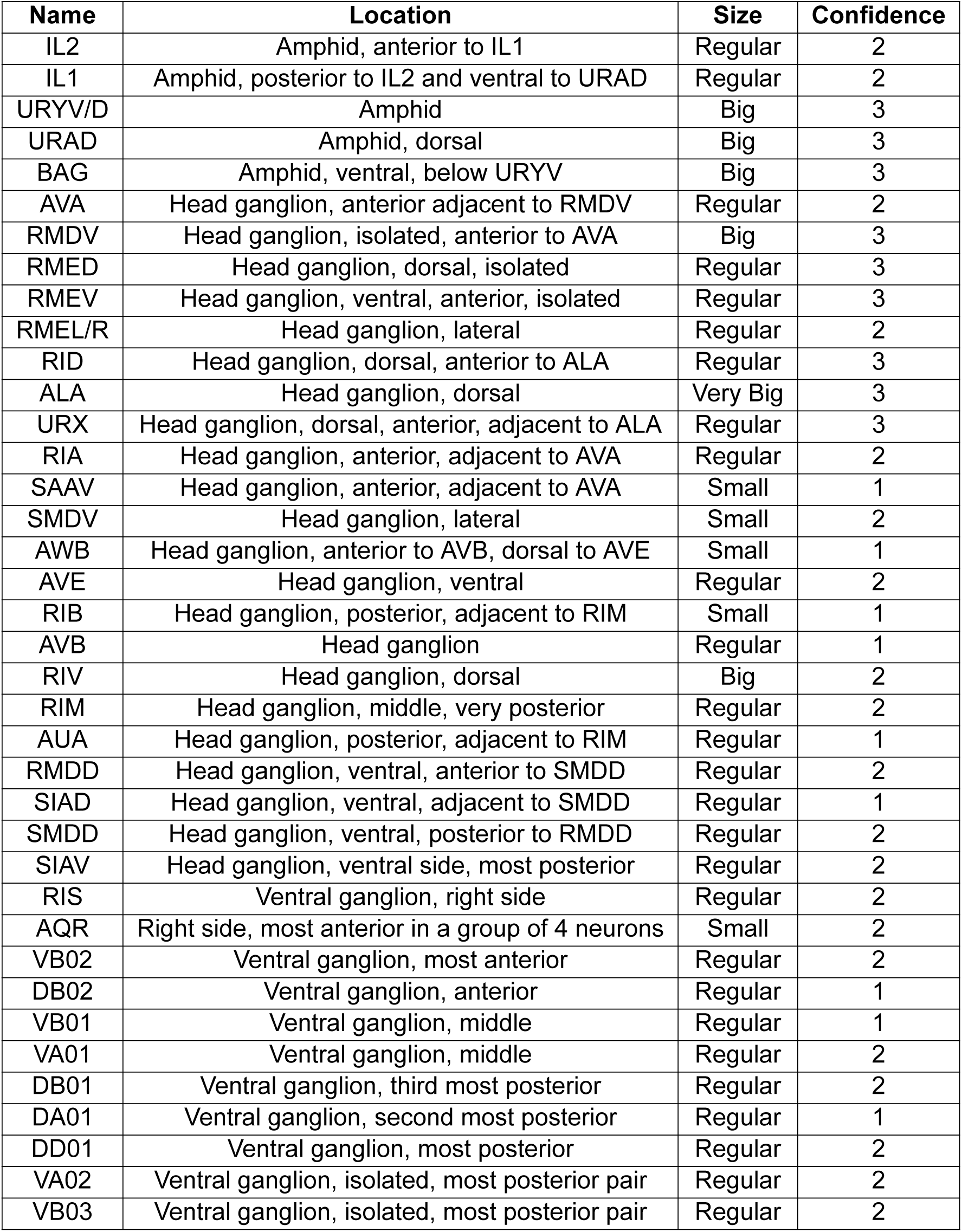
List of identified neuronal classes in whole-brain datasets. The anatomical region, general shape and confidence level of each neuronal class are indicated. Note that the anatomical shape is per the expression of the pan neuronal promoter we used (unc-31p) and may change due to different expression levels when using different promoters. Confidence levels range from 0 to a maximum of 3. Confidence level 3 indicates neuronal classes that can be identified solely by anatomy and shape, confidence level 2 indicates neuronal classes identified by their anatomy and their unique stereotypic forward/reverse-tied and ventral/dorsal-related oscillatory activity pattern. Confidence levels 1 indicate neurons for which either the activity pattern and position are not unique enough for fully confident identification, or that the neuronal trace quality from that region does not allow confident identification based on activity. Confidence levels are the same in freely moving and immobilized recordings.

**Table S4.** Description of the strains used in this study.

| Strain name | Genotype | Description | Relevant Figures |
| --- | --- | --- | --- |
| ZIM2165 | Mzmls133;lite-1(ce314) X. | Mzmls113=[unc-31p::SV40-NLS::jGCaMP7b::egl-13-NLS;unc-31p::SV40-NLS::mScarlet::egl-13-NLS] | All Figures |
| ZIM2498 | Mzmls113; lite-1 (ce314) X., gcy-35(ok769) I.; gcy-31(ok296) X., gcy-9(n4470) X. | Mzmls113=[unc-31p::SV40-NLS::jGCaMP7b::egl-13-NLS; unc-31p::SV40-NLS::mScarlet::egl-13-NLS] | Figs. 4 and S8 |
| ZIM2319 | mzmls119;lite-1(ce314) X. | Mzmls119 = [unc-31p::SV40-NLS::GFP::egl-13-NLS; unc-31p::SV40-NLS::mScarlet::egl-13-NLS] | Fig.3<br>Fig.2 and Fig.S4 |
| ZIM2521 | lite-1(ce314) X.;Mzmls113; Mzmls120 | Mzmls113=[unc-31p::SV40-NLS::jGCaMP7b::egl-13-NLS; unc-31p::SV40-NLS::mScarlet::egl-13-NLS],<br>Mzmls120=[mlc-2p::HisCl::rpl-3 3'UTR;flp-17p::mScarlet] | Fig. S8 |
| ZIM2474 | lite-1(ce314) X.;Mzmls52; otIs670 | Mzmls52= [unc-31p::SV40-NLS-GCaMP6f-egl-13-NLS]; | Methods |

